# Immunotherapy for ovarian cancer is improved by tumor-targeted delivery of a neoantigen surrogate

**DOI:** 10.1101/2023.10.11.561944

**Authors:** Lauren Rose Scanlon, Lisa Gabor, Olivia Khouri, Shahbaz Ahmad, Evan Levy, Elizabeth Buckley, Ariene Ouedraogo, Dennis Yi-Shin Kuo, Ken Lin, Nicole Nevadunsky, Sara Isani, Shixiang Sun, Claudia Gravekamp

**Affiliations:** Department of Microbiology and Immunology, Albert Einstein College of Medicine, Forchheimer Building, 1300 Morris Park Avenue, Bronx, NY 10461, USA; Montefiore Medical Center/Albert Einstein College of Medicine, Division of Gynecologic Oncology, 1695 Eastchester Road, Bronx, NY 10461, USA; Department of Genetics, 1300 Morris Park Avenue, Bronx, NY 10461, USA

**Keywords:** ovarian cancer, Listeria, tetanus toxoid, T cell responses, recall antigen, neoantigen surrogate

## Abstract

Ovarian cancer is known for its poor neoantigen expression and strong immunosuppression. Here, we utilized an attenuated non-pathogenic bacterium *Listeria monocytogenes* to deliver a highly immunogenic Tetanus Toxoid protein (Listeria-TT), as a neoantigen surrogate, into tumor cells through infection in a metastatic mouse ovarian cancer model (Id8p53-/-Luc). Gemcitabine (GEM) was added to reduce immune suppression. Listeria-TT+GEM treatments resulted in tumors expressing TT and reactivation of pre-existing CD4 and CD8 memory T cells to TT (generated early in life). These T cells were then attracted to the TT-expressing tumors now producing perforin and granzyme B. This correlated with a strong reduction in tumor burden, and significant improvement of the survival time compared to all control groups. Checkpoint inhibitors have little effect on ovarian cancer partly because of low neoantigen expression. Here we demonstrated that Listeria-TT+GEM+anti-PD1 was significantly more effective (efficacy and survival) than anti-PD1 or Listeria-TT+GEM alone. Of clinical interest, high doses of anti-PD1 (PD1H) (when added to Listeria-TT+GEM) were less effective than the low doses (PD1L). IHC and ELISPOT demonstrated that high doses of anti-PD1 inhibited T cell function in the TME. Using RNAseq, Differentially Expressed Genes (DEG) analysis and Genes Set Enrichment Analysis (GSEA) showed that gene expression levels and biological pathways were predominantly upregulated in the PD1H compared to the PD1L group, in correlation with low immune infiltration in tumors, more immune suppression, and more aggressive ovarian cancer. In summary, this study suggests that our approach may benefit ovarian cancer patients.

Graphical Abstract
**Human concept:** Childhood vaccinations with the highly immunogenic tetanus toxoid (TT) generate TT-specific memory T cells, which circulate in the blood for life. After appearance of ovarian cancer (late in life), the patients will receive one high dose with Listeria-TT to deliver TT into tumor cells, followed by multiple low doses of Listeria-TT over a period of 2 weeks to restimulate the pre-existing memory T cells to TT. MDSC are involved in the delivery of Listeria-TT to the TME. Reactivated memory T cells will in turn destroy the tumor cells expressing TT. Multiple low doses of GEM will be added after TT has been delivered at the tumor site, which reduce immune suppression by eliminating MDSC and TAM (not shown here). Since individuals have seen TT earlier in life (during childhood vaccinations) and since TT is highly immunogenic (attracting T cells to the TME) but not expressed in normal cells, TT functions here as a vaccine recall antigen and as a neoantigen surrogate, respectively.

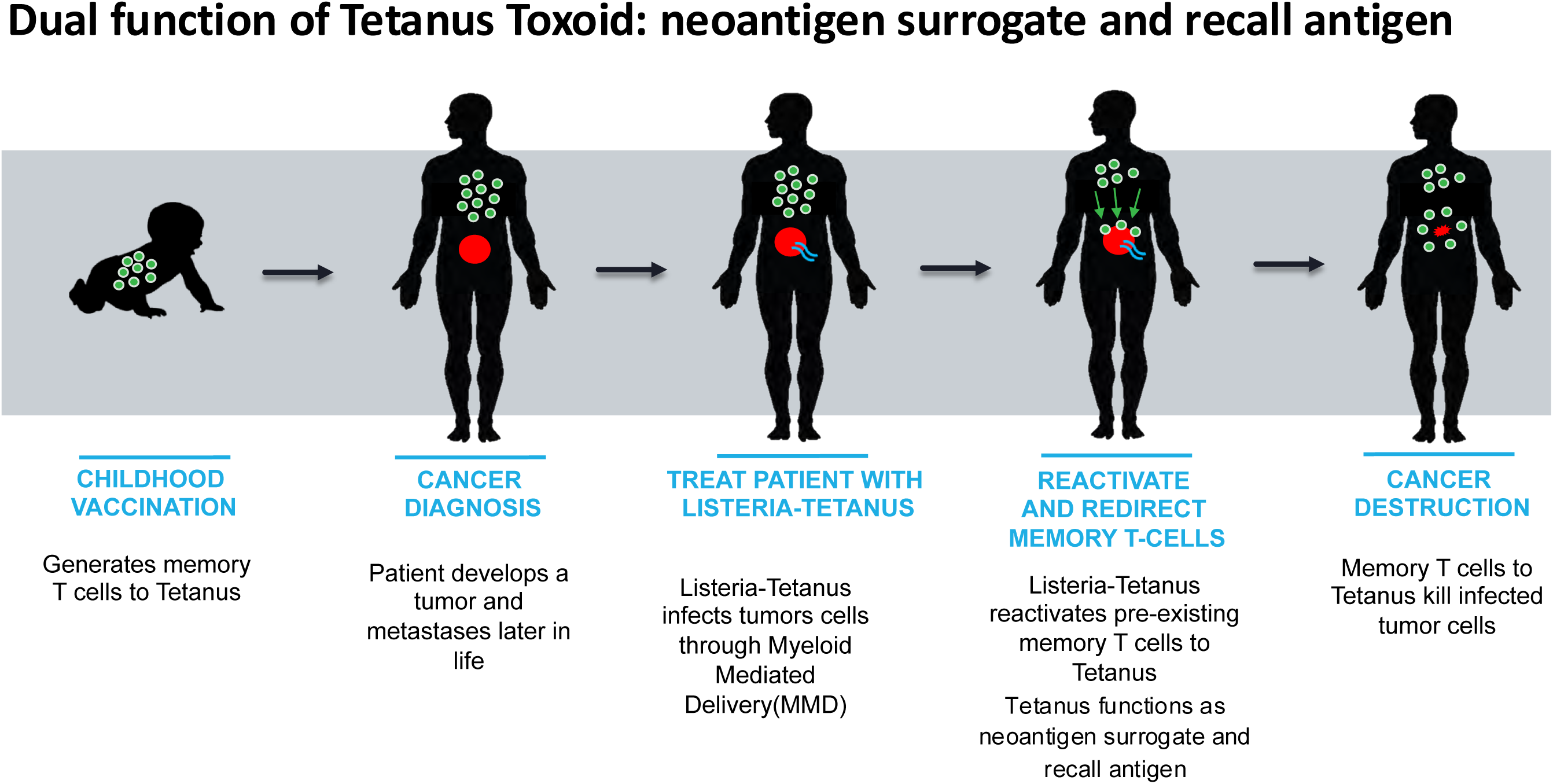

## Introduction

Globally, 225,500 new cases of ovarian cancer are diagnosed each year, with 140,200 cancer-specific deaths^1–3^. Ovarian cancer is a disease subdivided into different histological subtypes that have different risk factors which require different treatments^4^. Newly diagnosed cancer is treated with cytoreductive surgery and platinum-based chemotherapy. In recurrent cancer, chemotherapy, anti-angiogenic agents and poly ADP-ribose polymerase inhibitors are used. High grade-serous carcinoma (HGSC) is the most common histological subtype of ovarian cancer, that initially is highly responsive to platinum-based chemotherapy, but frequently relapses and becomes increasingly chemoresistant^5^. It has been reported that these relapses are caused by unlimited self-renewal and proliferative capacity of cancer stem cells (CSC)^6^. This underlines the desperate need for new alternative approaches.

While checkpoint inhibitors have shown impressive results for melanoma and lung cancer, this is less so for ovarian cancer^7^. This is partly because ovarian cancer has low levels of neoantigen expression^8^. We hypothesized that ovarian cancer patients would benefit from Listeria-TT. In a previous study, we showed that Listeria delivers highly immunogenic recall antigens, like tetanus toxoid (TT), to the tumor microenvironment (TME) and inside tumor cells of the poorly immunogenic pancreatic cancer^9^. Once injected, Listeria attracts and infects myeloid-derived suppressor cells MDSC^9–11^. These MDSC are also attracted to the cancer by chemo-attractants and cytokines^12,13^, and deliver the Listeria as a trojan horse to the TME, where they spread from MDSC into tumor cells, and from tumor cell to tumor cell, secreting TT protein^9^. Immune suppression in the TME allows Listeria to survive and multiply, but not in normal tissues that lack immune suppression^9^. Once Listeria has been settled in the TME, low doses of Gemcitabine (GEM) were added to reduce immune suppression in the TME. We found that high levels of TT in the TME attracted the pre-existing TT-specific memory CD4 T cells (generated earlier in life) to the TME, producing perforin and granzyme B, resulting in a strong reduction of tumors and metastases and significant longer survival. Thus, TT functions here as a recall antigen and as a neoantigen surrogate. Based on these results, the current study tested whether Listeria-TT+GEM was also effective against metastatic ovarian cancer. We hypothesized that delivery of neoantigen surrogate TT to the TME, would improve the efficacy of checkpoint inhibitors in mouse models of ovarian cancer.

Here, we showed that Listeria-TT+GEM is highly effective against metastases and tumors in the ovarian cancer Id8p53-/-Luc model (efficacy and survival), and that this correlated with a significant increase in CD4 and CD8 T cells producing perforin and granzyme B. Most interestingly, Listeria-TT+GEM+anti-PD1 antibodies were significantly more effective than anti-PD1 or Listeria-TT+GEM alone (efficacy and survival), and more treatment cycles significantly further improved the survival of mice compared to those that received one treatment cycle. Low doses of anti-PD1 were significantly more effective (efficacy and survival and CD4 T cell responses) than high doses when added to Listeria-TT+GEM. More detailed analysis (IHC and ELISPOT) revealed that T cell responses were reduced by high doses of anti-PD1, while analysis of gene expression profiles and biological pathways (RNAseq) showed lower immune cell infiltration, more immune suppression, more migration and invasion, in correlation with more aggressive ovarian cancer in the PD1H group. Potential mechanism(s) of improvement will be discussed. In conclusion, these results are promising for those ovarian cancer patients with low neoantigen expression.

## Results

### *Listeria*-TT infection of tumor cells *in vitro*

In previous studies we have shown that Listeria infects human and mouse pancreatic and breast tumor cells ^9,11,14,15^. Here, we showed that Listeria also infects mouse and chemoresistant human ovarian cancer cells *in vitro*. The infection rate of Id8p53-/-(mouse) or Hey (human) tumor cells was determined by culturing the tumor cells with *Listeria*-TT for 1hr, followed by lysis in water, and plating the lysate on agar. The number of CFU of Listeria-TT was counted the following day (figure S1A).

Previously, we showed that Listeria destroys human (MCF7) and mouse (4T1) breast tumor cells through the activation of the NADPH-oxidase pathway, resulting in mitochondrial reactive oxygen species and subsequent tumor cell death ^14^. Here, we demonstrated that *Listeria*-TT also efficiently kills Id8 and Hey tumor cells *in vitro* **(figure S1BC).** For this purpose, tumor cells were incubated for 2 hours with Listeria-TT, followed by gentamicin, and then were cultured overnight. Dead and alive tumor cells were counted based on trypan blue staining.

### The Id8p53-/- Luc model

p53 is mutated in 80% of high grade serous ovarian cancers. Therefore, we used an ovarian cancer model with deletions in p53 and expression of Luciferase (Id8p53-/- Luc model)^16,17^. The expression of Luciferase allowed us to analyze real-time effect(s) of therapy in live mice with ovarian cancer. We generated two models: the peritoneal model and the ovarian model. Using the peritoneal model, tumor cells were directly injected into the peritoneal cavity resulting in metastatic tumors dispersed throughout the peritoneal cavity (on the surface of the liver, diaphragm, pancreas, and gastrointestinal (GI) tract) similar to high-grade serous ovarian cancer **(figure S2A).** However, no tumor cells were found inside the ovaries **(figure S2B).** Therefore, we generated an ovarian model in which we injected tumor cells directly into the bursa of the ovaries, resulting in big ovarian tumors that later metastasized to the peritoneal cavity as well **(figure S2CD).**

### Accumulation of Listeria-TT in the TME

Before testing Listeria-TT as immunotherapy against ovarian cancer we first determined whether Listeria-TT accumulated in the TME. Briefly, 2x10^6^ Id8p53-/-tumor cells were injected intraperitoneally (IP). Four weeks later ascites was drained, followed by the IP injection of one high dose of Listeria-TT (10^7^ CFU). Next, bacteria were isolated from all metastases and normal tissues on both day 1 and 3 after injection of Listeria-TT, and the number of CFU of the bacteria per gram tissue was calculated. We found that Listeria-TT predominantly accumulated in the metastases, with the highest number in the metastases of the diaphragm, GI and liver (**figure 1A**).

**Figure 1:**
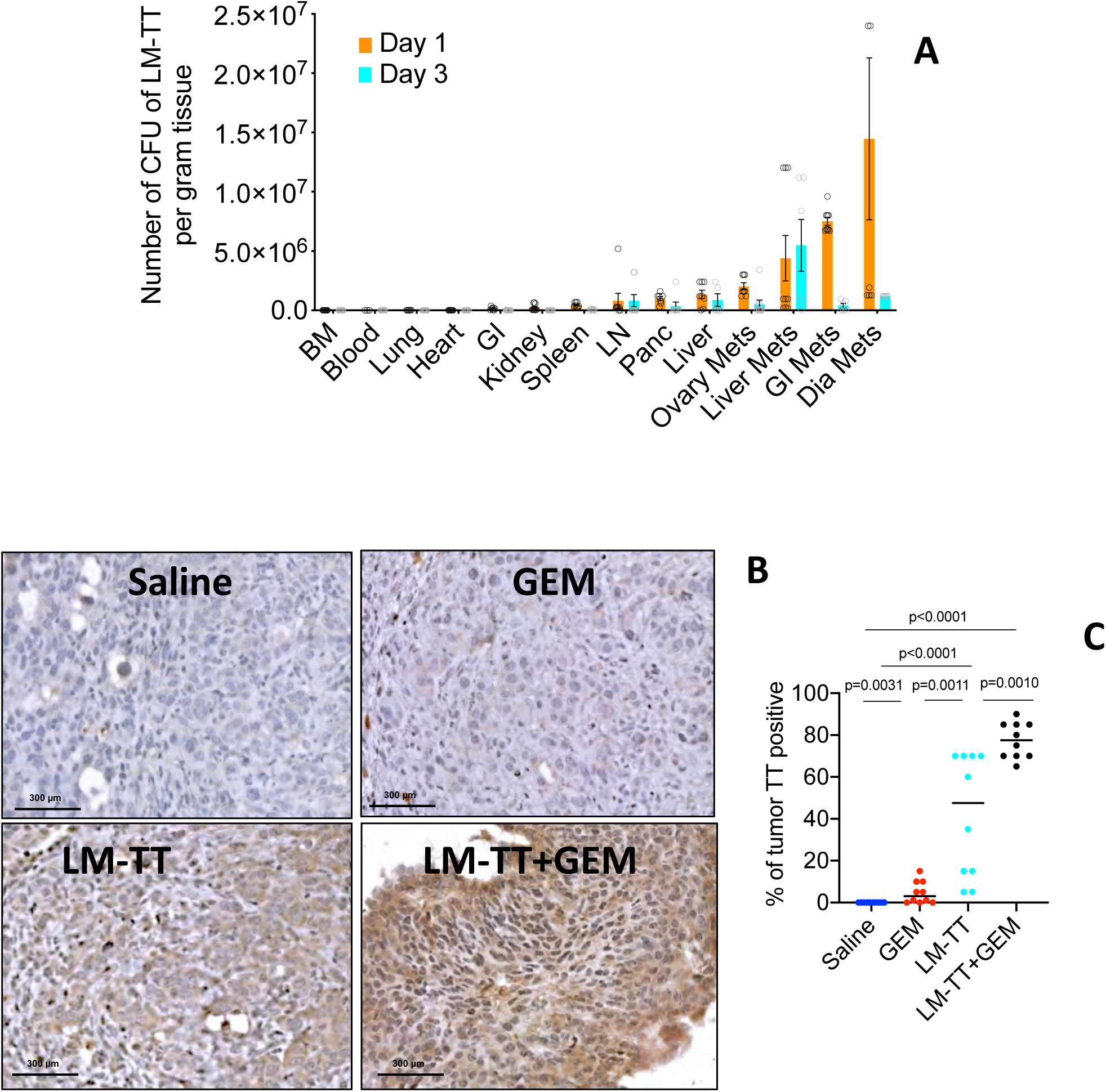
Listeria-TT accumulates and delivers TT in the TME. **(A)** Biodistribution of Listeria-TT in the ovarian cancer model Id8p53-/- (peritoneal model). Listeria-TT accumulated in the metastatic tumors of the Id8p53-/-Luc (peritoneal model). n=3 mice per time point. From each tissue 3 pieces were analyzed in triplicate. The number of CFU of Listeria-TT was determined per gram tissue or per mL blood. **(B)** Listeria delivers TT to the TME (ovarian cancer model) as shown by IHC, and quantified **(C).** 10 fields in each group were counted and averaged. n=3 mice per group. Statistical significance was determined by using Mann-Whitney, p<0.05 is significant. Error bars represent standard error of the mean (SEM). LM-TT=Listeria-TT. GEM=Gemcitabine.

### *Listeria*-TT+GEM attracts CD4 and CD8 T cells to the TME producing perforin and Granzyme B

Here, we confirmed that Listeria delivered TT to the ovarian tumors (ovarian model). For this purpose, we treated the mice with a complete cycle of Listeria-TT+GEM (pre-vaccinations and Listeria-TT+GEM) **(figure S3A)** and analyzed the tumors by immunohistochemistry (IHC) for the expression of TT. As shown in **figure 1BC**, TT was expressed in 50% and 80% of the tumor area of mice that received Listeria-TT or Listeria-TT+GEM, respectively. Ovarian tumors are immunologically cold with few T cells present in the TME of mice or humans ^18^. After a complete treatment cycle of the Id8p53-/-Luc (ovarian model) with Listeria-TT+GEM, we found that the number of both CD4 and CD8 T cells significantly increased compared to the saline group (**figure 2AB and figure S4AB).** This differs in comparison to the pancreatic tumors of KPC mice treated with a similar therapy regimen, in which we observed CD4 T cells only ^9^. Also, in the GEM-treated mice the number of CD4 and CD8 T cells was significantly increased compared to the saline group. This is not surprisingly since GEM eliminates the immune suppressive myeloid-derived suppressor cells (MDSC) and tumor-associated macrophages in the TME and in blood ^9^, which negatively affect T cell trafficking ^19^.

**Figure 2:**
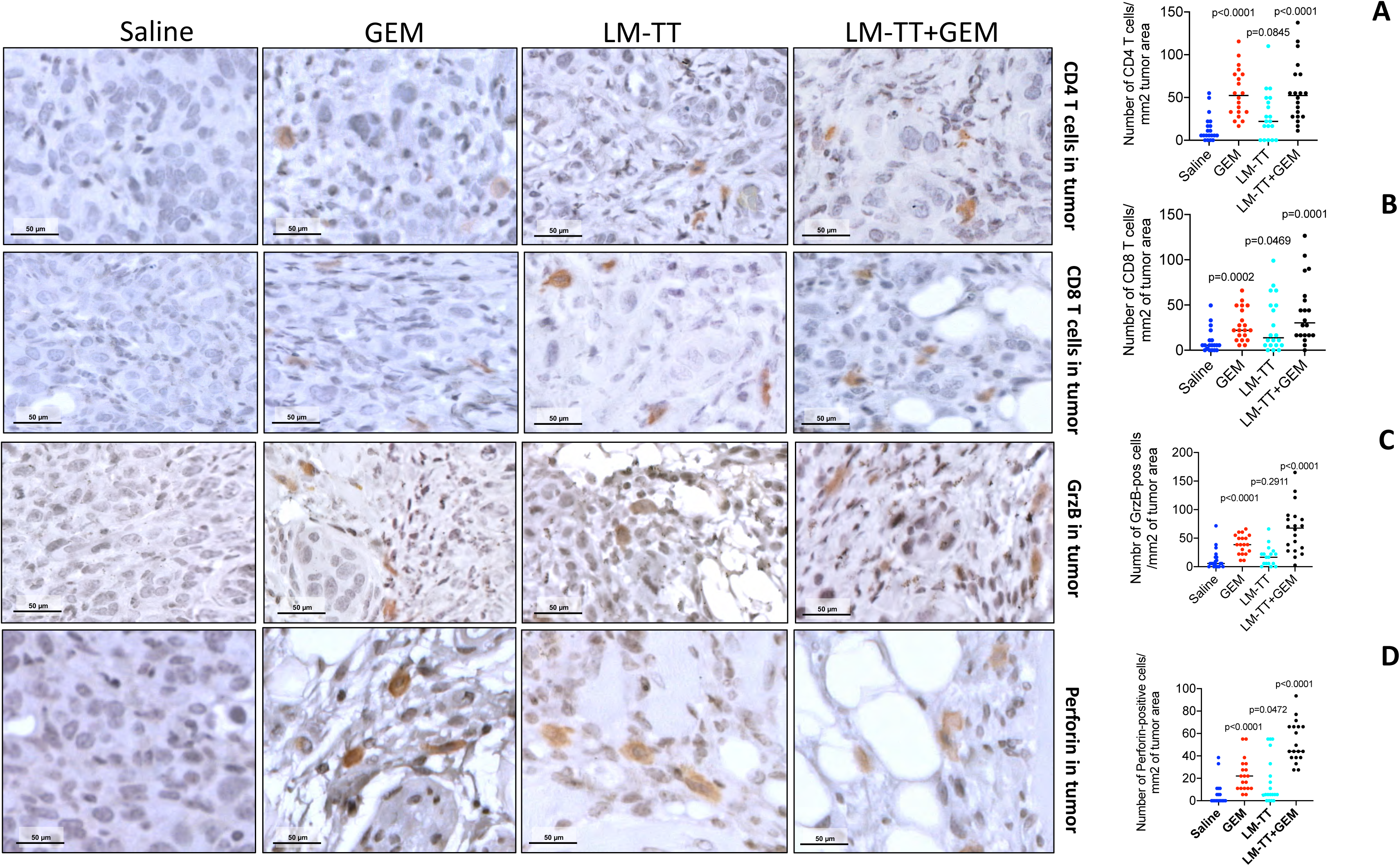
CD4 and CD8 T cells are attracted to and activated in the ovarian tumors by Listeria-TT+GEM treatment in the Id8p53-/- Luc model. The number of CD4 **(A)** and CD8 **(B)** T cells increased in the TME after a complete treatment cycle with Listeria-TT+GEM treatment in Id8p53-/- ovarian model. Listeria-TT+GEM increased the production of Granzyme B **(C)** and Perforin **(D)** in the Id8p53-/- ovarian tumors. 20 fields per group were counted and the results were averaged. The average number of positive cells were calculated per square millimeter tumor area. n=3 mice per group. Statistical significance was determined by using Mann-Whitney, p<0.05 is significant. All groups in **fig A-D** were compared to the Saline group. Error bars represent SEM.

To show that these T cells were functional, we also analyzed the production of perforin and granzyme B in the TME by IHC. We found that the production of perforin and granzyme B was significantly increased in mice treated with LM-TT+GEM, compared to all control groups (**figure 2CD and figure S4CD).** Of note, MDSC and tumor-associated macrophages (TAM) strongly inhibits T cell function through different pathways ^9,20^. Previously, we have shown that GEM eliminate these MDSC and TAM ^9^.

### *Listeria*-TT+GEM reduces ovarian cancer in Id8p53-/-Luc mice (peritoneal model) and improves survival

After evaluating the effect of *Listeria*-TT+GEM on immune responses in the TME, we also analyzed the therapeutic effect on ovarian cancer in the Id8p53-/- Luc mice. For this purpose, we generated TT-specific memory T cells in C57BL/6 mice prior to tumor development (6-8 weeks of age) by injecting the TT vaccine im twice one week apart (the same vaccine be given to humans). One week after the last vaccination, Id8p53-/-Luc tumor cells were injected intraperitoneally (peritoneal model). After the presence of metastases were confirmed by H&E and IVIS (**figure 3AB**), the mice were treated with *Listeria*-TT+GEM as outlined in **figure S3A.** The region of interest (ROI) of the luciferase signal (measurement for tumor growth) was determined by IVIS before treatment (day 0), during treatment (day 10) and after treatment (day 20). On day 20, the IVIS signal was significantly lower in the Listeria-TT+GEM compared to all other groups (**figure 3BC**). After the last treatment, the mice were monitored without any further treatment in a survival study. Confirming the IVIS data, mice treated with Listeria-TT+GEM lived significantly longer than the mice in all other treatment groups (**figure 3D**).

**Figure 3:**
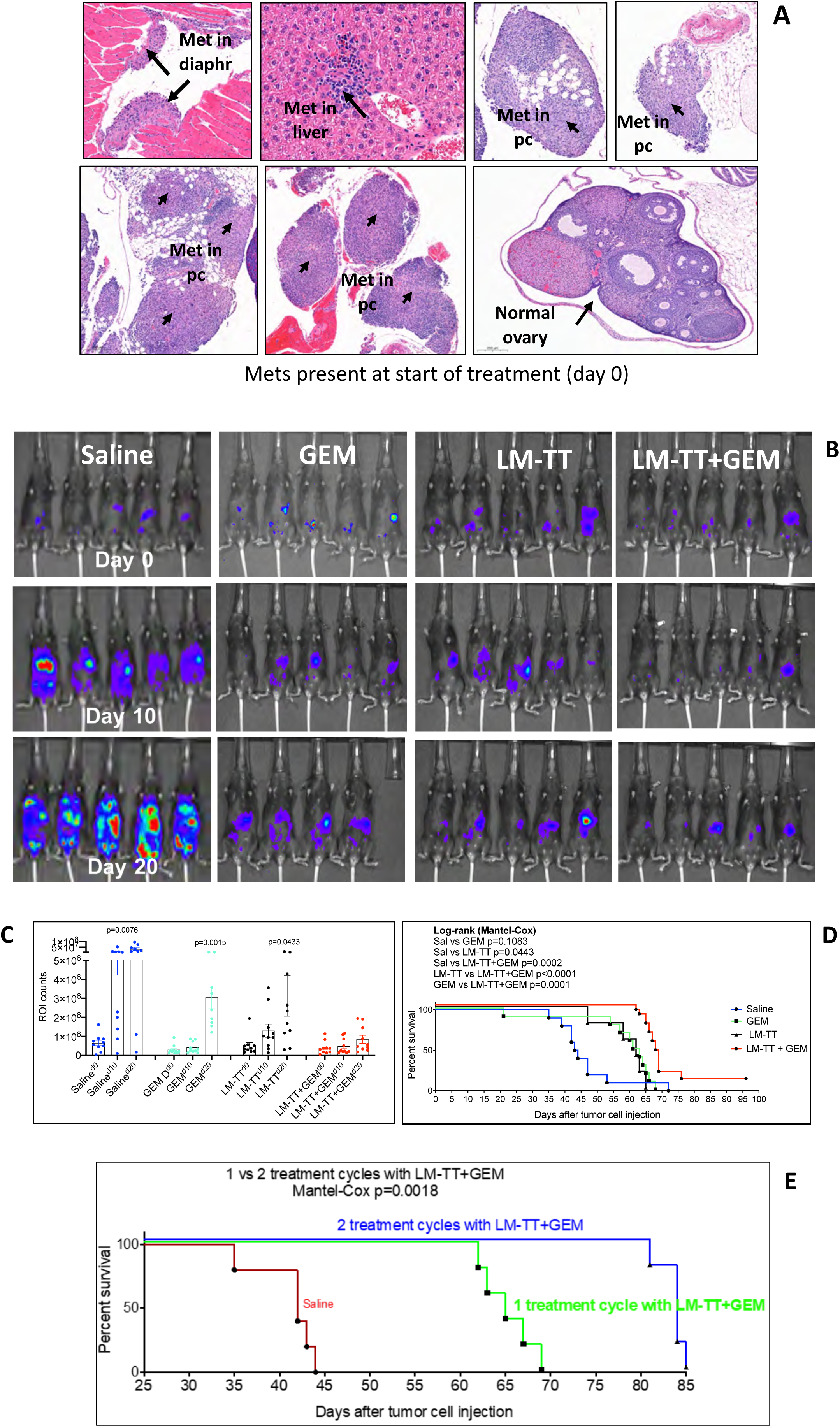
Listeria-TT+GEM significantly inhibits growth of ovarian metastases and improves survival time compared to all control groups in the Id8p53-/- Luc peritoneal model. At the start of treatment, metastases are visible in the peritoneal cavity (in diaphragm, liver and along GI) by H&E staining **(A)** and by IVIS **(B).** Listeria-TT+GEM significantly inhibits the growth of metastatic tumors in the peritoneal cavity (pc) as shown by IVIS **(B)** and by ROI counts quantified **(C).** All groups were compared to the LM-TT+GEM group at day 20 of treatment. Listeria-TT+GEM significantly improves the survival of ID8p53-/- Luc mice compared to all control groups **(D). In fig C and D,** n=10 mice per group. Two treatment cycles of Listeria-TT+GEM significantly improved the survival time compared to one treatment cycle or saline **(E).** n=5 mice per group. Statistical significance was determined for efficacy by using Mann-Whitney, p<0.05 is significant, and for survival by using Mantel-Cox, p<0.05 is significant. Error bars represent SEM.

We observed that TT was expressed in 80% of the ovarian tumors (**figure 1BC**). This indicates that 20% of the primary tumor does not express TT, which will grow back because only TT-positive tumor cells will be destroyed by the TT-specific memory T cells. To address this problem, we followed up with a second treatment cycle aiming to deliver TT to the TT-negative tumor cells. As shown in **figure 3E**, we increased the survival time by 50% after two treatment cycles compared to untreated mice, and by 23% compared to one treatment cycle.

### *Listeria*-TT+GEM reduces ovarian cancer in Id8p53-/-Luc mice (ovarian model) and improves survival

A similar treatment protocol of Listeria-TT+GEM has been evaluated in the ovarian model **(figure S3A).** Primary tumors developed 11 days after injection of tumor cells into the ovaries. Tumor growth was determined by tumor weight (at the end of treatment) (**figure 4A**) or by IVIS (days 27, 37, and 47) (**figure 4B**). The IVIS signal was significantly reduced in the Listeria-TT+GEM group compared to the saline group and reduced 2-fold over time (before vs after treatment) within the group that received Listeria-TT+GEM (**figure 4C**). However, little effect of GEM or Listeria-TT was observed on the primary tumors. Survival data confirmed the results by tumor weight and IVIS signal, i.e. mice treated with Listeria-TT+GEM lived significantly longer than all other groups (**figure 4D**). Of note, the tumor cells multiplied much more abundantly in the peritoneal model than in the ovarian model. Also, the 50% survival time of untreated mice with ovarian tumors was 62 days, while for mice with the peritoneal metastases this was only 44 days.

**Figure 4:**
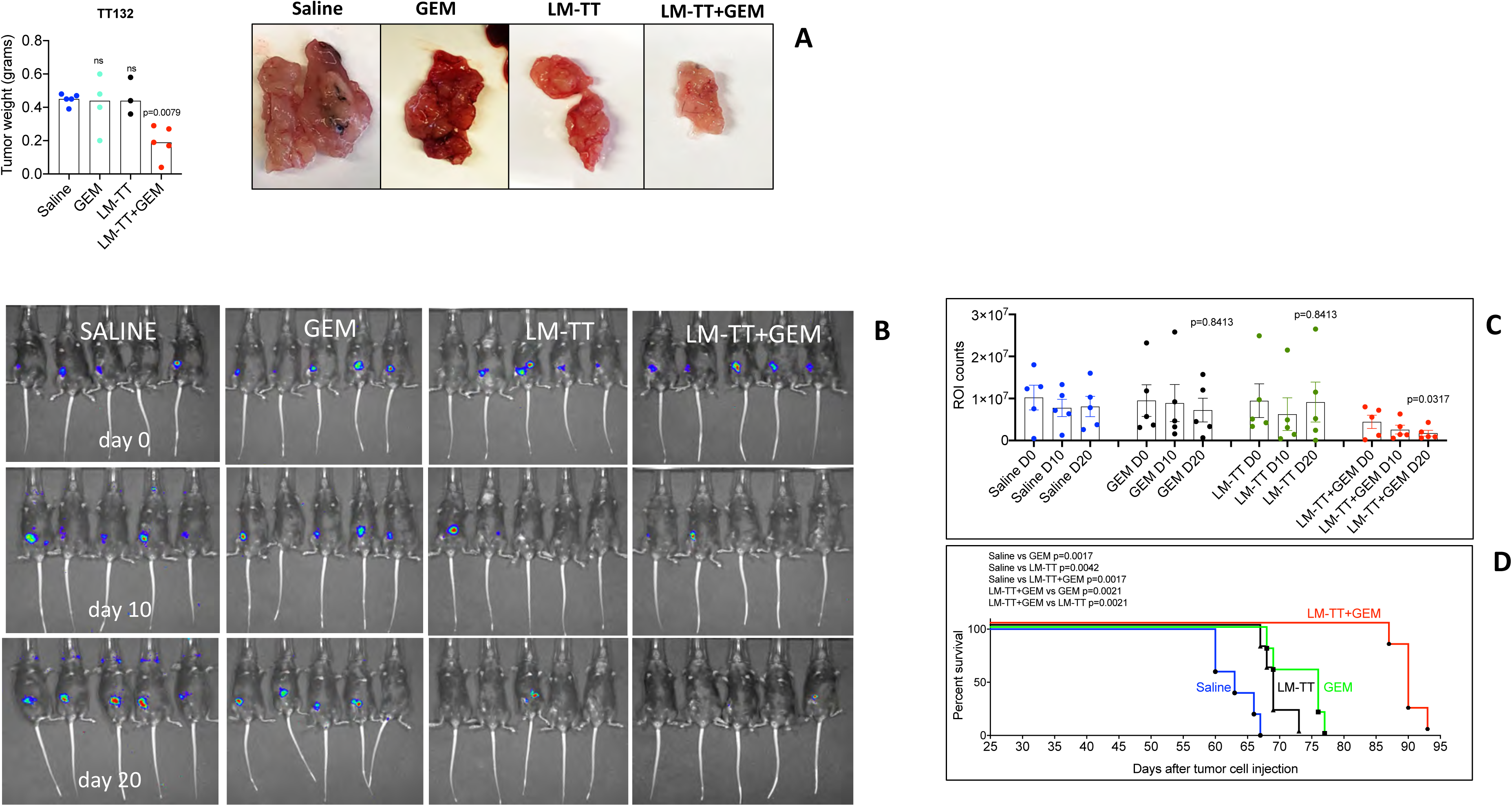
Listeria-TT+GEM significantly reduces ovarian tumors and improves the survival time of the Id8p53-/- Luc ovarian model. Listeria-TT+GEM significantly reduces the weight of the ovarian tumors **(A).** In this experiment, mice were euthanized 4 weeks after the last treatment. In a second experiment, ovarian tumors were analyzed by ROI counts by IVIS compared to the saline group **(BC)** on days 0, 10 and 20 of treatment. All groups were compared to the Saline group at day 20 of the treatment. Listeria-TT+GEM significantly improves the survival time of Id8p53-/-Luc mice compared to all control groups **(D).** n=5 mice per group. Statistical significance was determined for efficacy by using Mann-Whitney, p<0.05 is significant, and for survival by using Mantel-Cox, p<0.05 is significant. Error bars represent SEM.

### Listeria-TT+GEM+anti-PD1 is significantly more effective than anti-PD1 or Listeria-TT+GEM alone (peritoneal model)

In mice ^7,8^ and patients (NCT02580058, NCT02718417, NCT02674061) anti-PD1 has no effect on HGSC. This is partly because HGSC has poor neoantigen expression ^21^. One of our hypotheses is that delivery of TT as a neoantigen surrogate to the TME may improve the effect of checkpoint inhibitors. To prove our hypothesis, we delivered neoantigen surrogate TT by Listeria to the TME. More specifically, we combined the Listeria-TT+GEM treatment with anti-PD1 antibodies, and compared their effect to Listeria-TT+GEM or anti-PD1 alone treatments. Prior to these experiments, we first tested two different doses of anti-PD1 antibodies: a high dose of 130 μg/dose and a low dose 50 μg/dose. In the literature higher doses (130 μg/dose) were mostly used ^22,23^, but one report described Listeria-E7 combined with a low dose of anti-PD1 ^24^. Therefore, we tested both doses. For a schematic view of the treatment protocol see **figure S3B.** It appeared that the low dose anti-PD1 (50 μg/dose), was more effective than the high dose (**figure 5AB**), which was confirmed by a survival study (**figure 5C**). Subsequently, we tested whether GEM was required for optimal treatment. As shown in **figure 5DE**, Listeria-TT+GEM+anti-PD1 was more effective than Listeria-TT+anti-PD1 against ovarian metastases. We then tested Listeria-TT+GEM+anti-PD1 in comparison to Listeria-TT+GEM or anti-PD1 alone or the isotype control. While anti-PD1 alone showed an insignificant effect on the metastases in the Id8p53-/- model (**figure 5FG**) compared to the saline group, Listeria-TT+GEM+anti-PD1 was significantly more effective than anti-PD1 or Listeria-TT-GEM alone (**figure 5FG**). We used an isotype control for PD1, and as expected, the isotype control for PD1 + Listeria-TT+GEM showed similar results as Listeria-TT+GEM alone (**figure 5FG**). Also, in a survival study we showed that Listeria-TT+GEM+anti-PD1 was significantly more effective than all control groups (**figure 5H**).

**Figure 5:**
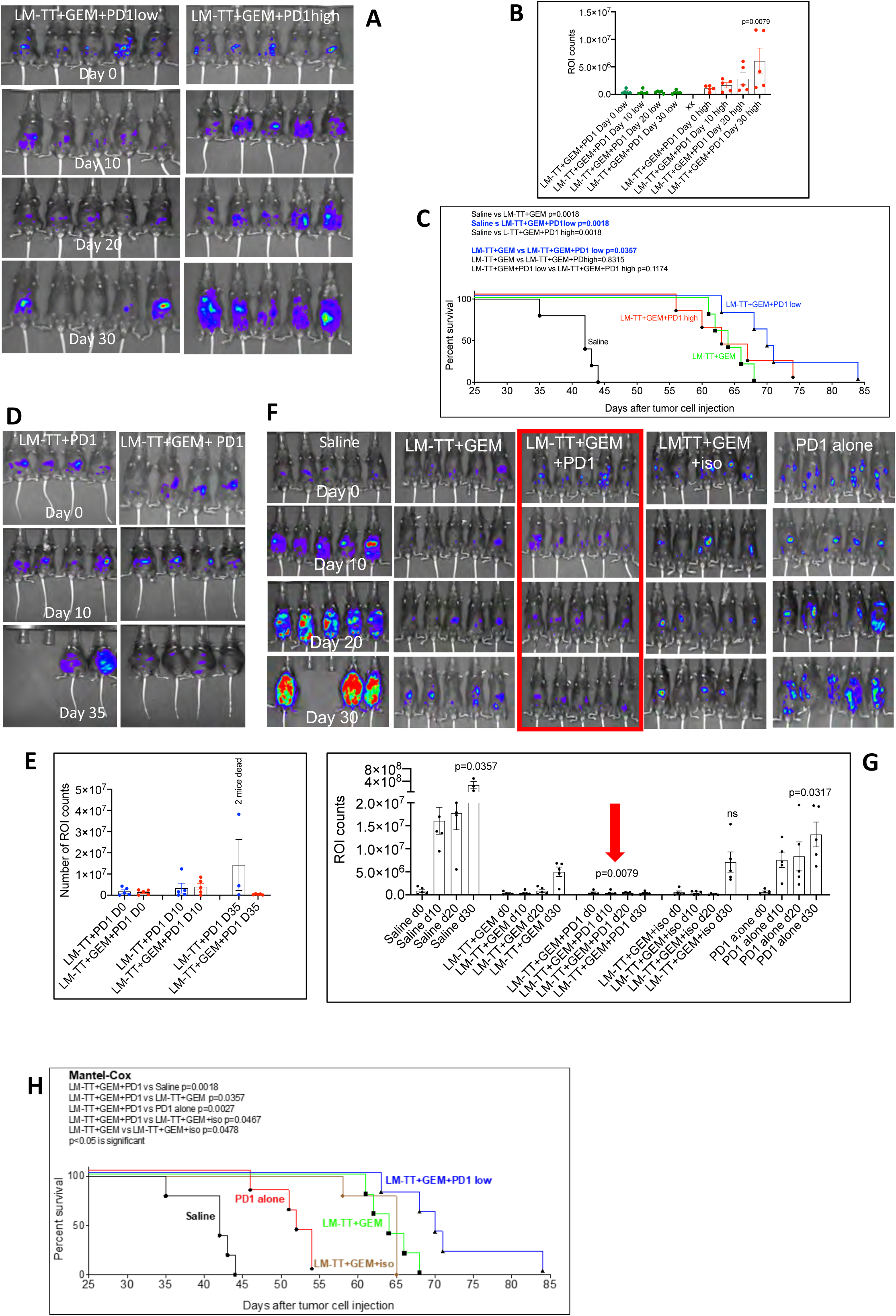
Low dose anti-PD1 is more effective than high dose anti-PD1 when combined with Listeria-TT+GEM (peritoneal model). Low dose anti-PD1 is more effective than high dose anti-PD1 when combined with Listeria-TT+GEM as shown by IVIS **(AB)** and in a survival curve **(C).** n=5 mice per group. GEM is required for an optimal effect of Listeria-TT+GEM+anti-PD1 in the ID8p53-/- peritoneal model **(DE).** n=4 mice per group. Mice treated with Listeria-TT+GEM+anti-PD1low has significantly less metastases compared to Listeria-TT+GEM, as shown by IVIS **(FG)** and significantly live longer compared to all control groups as shown by the survival curve **(H).** In fig **G,** all groups were compared to LM-TT+GEM at day 30 of treatment. n=5 mice per group. Statistical significance was determined for efficacy by using Mann-Whitney, p<0.05 is significant, and for survival by using Mantel-Cox, p<0.05 is significant. Error bars represent SEM.

To determine if we were able to further increase the efficacy we administered three treatment cycles with Listeria-TT+GEM+anti-PD1. As shown in **figure 6AB**, both Listeria-TT+GEM+anti-PD1 (3 treatment cycles) and Listeria-TT+GEM (3 treatment cycles) are significantly more effective against ovarian metastases than the saline on day 30 of treatment. On day 54, all saline-treated mice were dead, on day 60 all anti-PD1-treated mice, on day 91 all Listeria-TT+GEM-treated mice, while all Listeria-TT+GEM+PD1 mice were dead on day 120. This was confirmed by survival data (**figure 6C**). In **figure 6D**, we show that the survival time of mice that received three treatment cycles with LM-TT+GEM+anti-PD1 was significantly longer than those who received one treatment cycle or saline.

**Figure 6:**
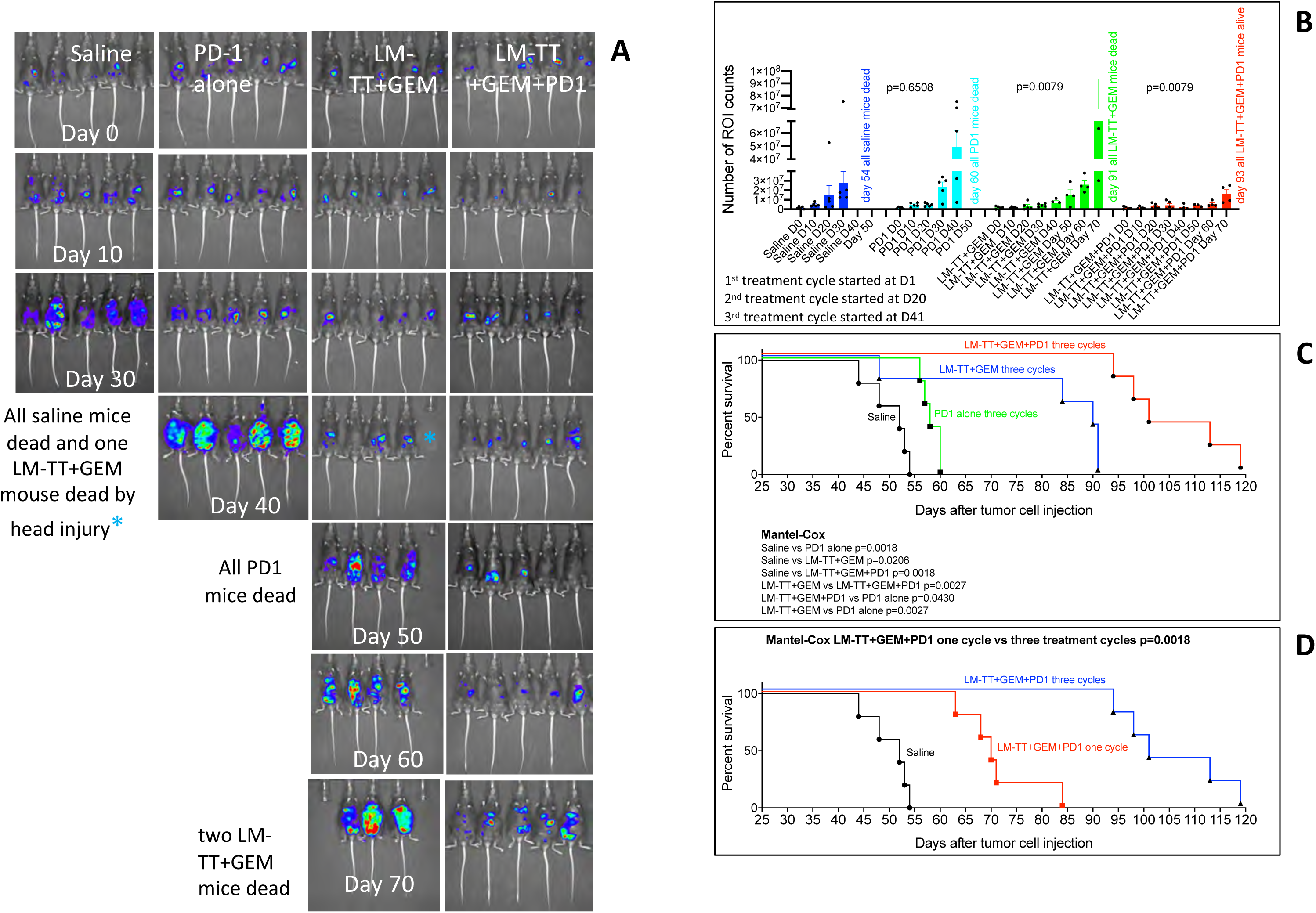
Three treatment cycles of Listeria-TT+GEM+anti-PD1 significantly further improves survival compared to one treatment cycle. Metastases were generated in the Id8p53-/-Luc peritoneal model. Both Listeria-TT+GEM+anti-PD1 and Listeria-TT+GEM were significantly more effective against ovarian metastases than the Saline group on day 30 of treatment, as shown by IVIS **(A),** and ROI counts quantified **(B).** Mice treated with LM-TT+GEM+anti-PD1 significantly lived longer than all control groups **(C).** We also compared three with one treatment cycle of LM-TT+GEM+anti-PD1 and found that mice receiving three treatment cycles significantly live longer compared to one treatment cycle **(D).** In fig D, mice with one and mice with three treatment cycles were performed as two independent experiments. LM-TT+GEM+anti-PD1 one cycle: 1x10E6 tumor cells injected, start treatment 11 days after tumor cell injection. LM-TT+GEM+anti-PD1 three cycles: 2x10E6 tumor cells injected, start treatment 14 days after tumor cell injection. Statistical significance was determined for efficacy by using Mann-Whitney, p<0.05 is significant, and for survival by using Mantel-Cox, p<0.05 is significant. Error bars represent SEM.

### Mechanistic mode of action of anti-PD1 when combined with Listeria-TT+GEM

It has been reported that Listeria upregulates PD-L1 on the surface of mouse dendritic cells in the spleen of the TC-1 model^24^ which may improve the effect of anti-PD1. We determined by IHC if Listeria also upregulated PD-L1 in the spleen of the Id8p53-/-Luc model after a full treatment cycle by counting the number of PD-L1-positive cells per square millimeter of spleen area. While the number of PD-L1-positive cells in spleens of mice treated with GEM or LM-TT+GEM was significantly higher than the saline group (p<0.0001), this effect was less robust and not significant for Listeria-TT alone (**figure 7AB**). Moreover, the saline group was almost completely negative for PD-L1 expression. Using the same mice we also determined the effect of Listeria-TT and/or GEM on PD-L1 expression in the ovarian tumors by IHC. Also, here the number of PD-L1-positive cells per square millimeter of tumor area was counted in all four treatment groups. Only GEM significantly upregulated PD-L1 in the primary tumors (p=0.0002) (**figure 7CD**).

**Figure 7:**
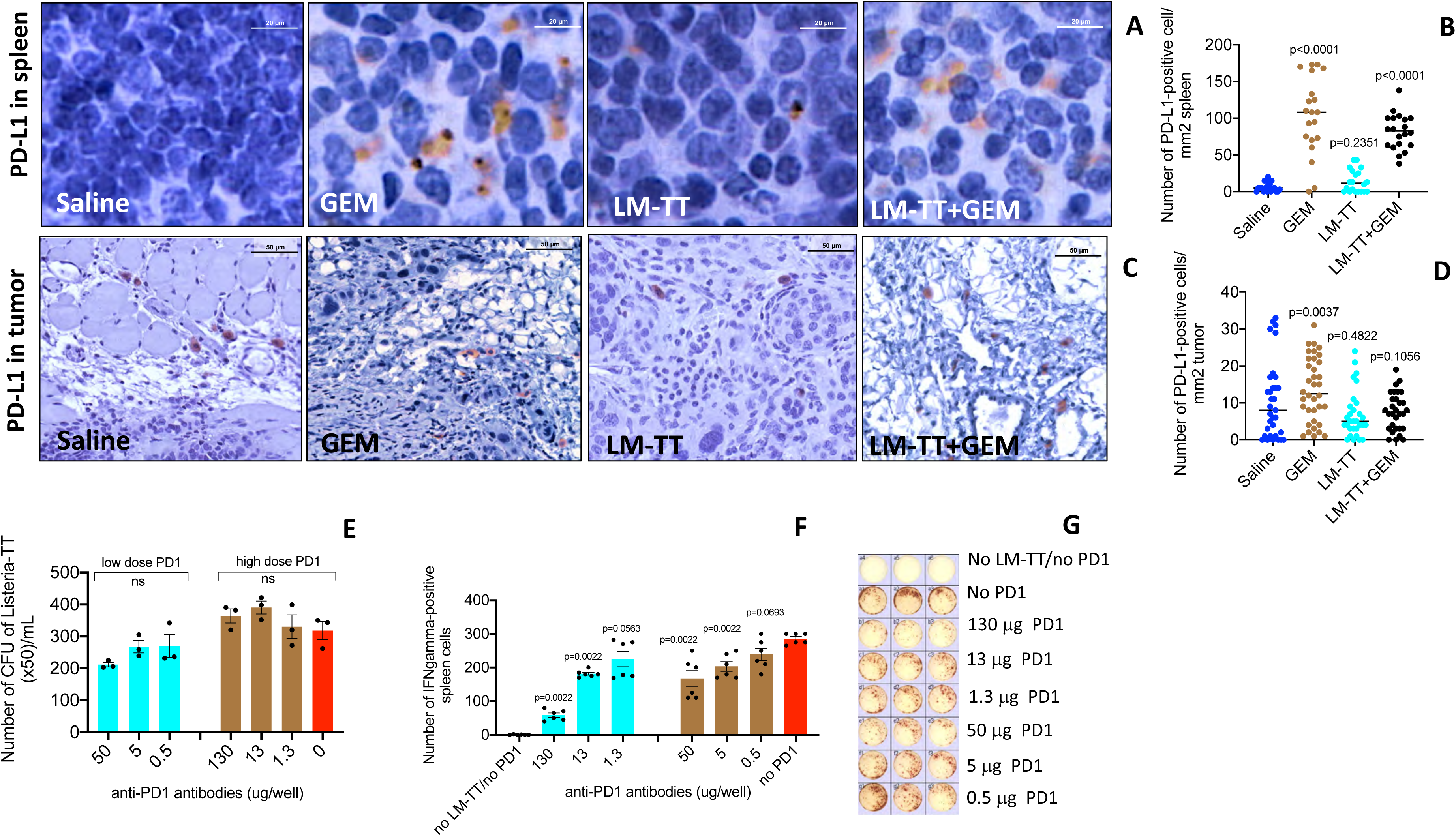
Mechanism of action of anti-PD1 antibodies *in vitro* and *in vivo*. IHC shows that GEM or LM-TT+GEM treatment but not LM-TT alone significantly upregulates PD-L1 expression in spleens *in vivo* **(AB)** of Id8p53-/- mice, while in ovarian tumors PD-L1 expression is upregulated by GEM only **(CD)**. All groups were compared to the Saline group. In fig A-D, n=3 mice per group, and 4 sections with 10 fields per section were analyzed. The results were averaged and analyzed by Mann-Whitney, p<0.05 is significant. Anti-PD1 does not affect the viability of Listeria-TT *in vitro* **(E).** High doses of anti-PD1 reduces the immune response to Listeria-TT (production of IFNγ) in the spleen of TT-vaccinated mice *in vitro* in an ELISPOT **(FG).** All groups were compared to immune cells lacking anti-PD1 antibodies (red bar) (with Listeria-TT infection). Results were analyzed by Mann-Whitney, p<0.05 is significant. Error bars represent SEM.

As shown in **figure 5**, low doses of anti-PD1 antibodies were more effective against ovarian metastases (efficacy (ROI) and survival) than high doses, when combined with Listeria-TT+GEM. To explain this difference in efficacy, we analyzed potential pathological effect(s) of high doses of anti-PD-1 (130 μg/dose) on normal tissues (liver, lungs, kidneys and GI tract) in comparison to those that received low doses (50 μg/dose) of anti-PD1. Minimal effects of the high or low doses anti-PD1 antibodies was observed on spleen, liver, kidneys, lungs and GI tract **(Table S1).** In conclusion, no pathological differences were observed between the high and low doses of anti-PD1.

Another possibility is that high doses of anti-PD1 reduce the viability of Listeria-TT. We therefore incubated Listeria-TT overnight with serial dilutions of high and low doses of anti-PD1 and plated them on LB agar. No significant difference in the number of CFU of Listeria-TT was observed between of high (130, 13, 1.3 and 0 μg/200 μL) or low (50, 5, 0.5 and 0 μg/200μL) doses of anti-PD1 antibodies (**Fig. 7E**), indicating that anti-PD1 had no effect on the viability of Listeria-TT *in vitro*.

Finally, we tested whether high doses of anti-PD1 affected the production of IFNγ of spleen cells (TT-specific T cells, macrophages) *in vitro.* For this purpose, mice were immunized twice with TTvacc one week apart. One week after the last immunization, 200,000 spleen cells were restimulated with Listeria-TT in an ELISPOT for 2 hrs, followed by gentamycin treatment during the restimulation period (to kill all extracellular Listeria), in the presence of serial dilutions of high and low doses of anti-PD1 antibodies as described above. Here we found an inverted dose-response effect of anti-PD1. Higher doses of anti-PD1 correlated with less production of IFNγ by spleen cells (**figure 7FG**). In other words, the function of immune cells was decreased by the high dose of anti-PD1 compared to low or no anti-PD1.

To analyze the function of T cells in more detail we also performed IHC on the ovarian tumors of mice treated with anti-PD1low or anti-PD1high in combination with Listeria-TT+GEM (**figure 8**). Here we showed that the number of CD4 T cells in the mice treated with anti-PD1H+Listeria-TT+GEM was significantly lower than in those treated with anti-PD1L+Listeria-TT+GEM, with significant lower production of Perforin and Granzyme B, suggesting that the T cells in the anti-PD1H group were not or less functional. The number of CD8 T cells were not affected by high compared to the low anti-PD1.

**Figure 8:**
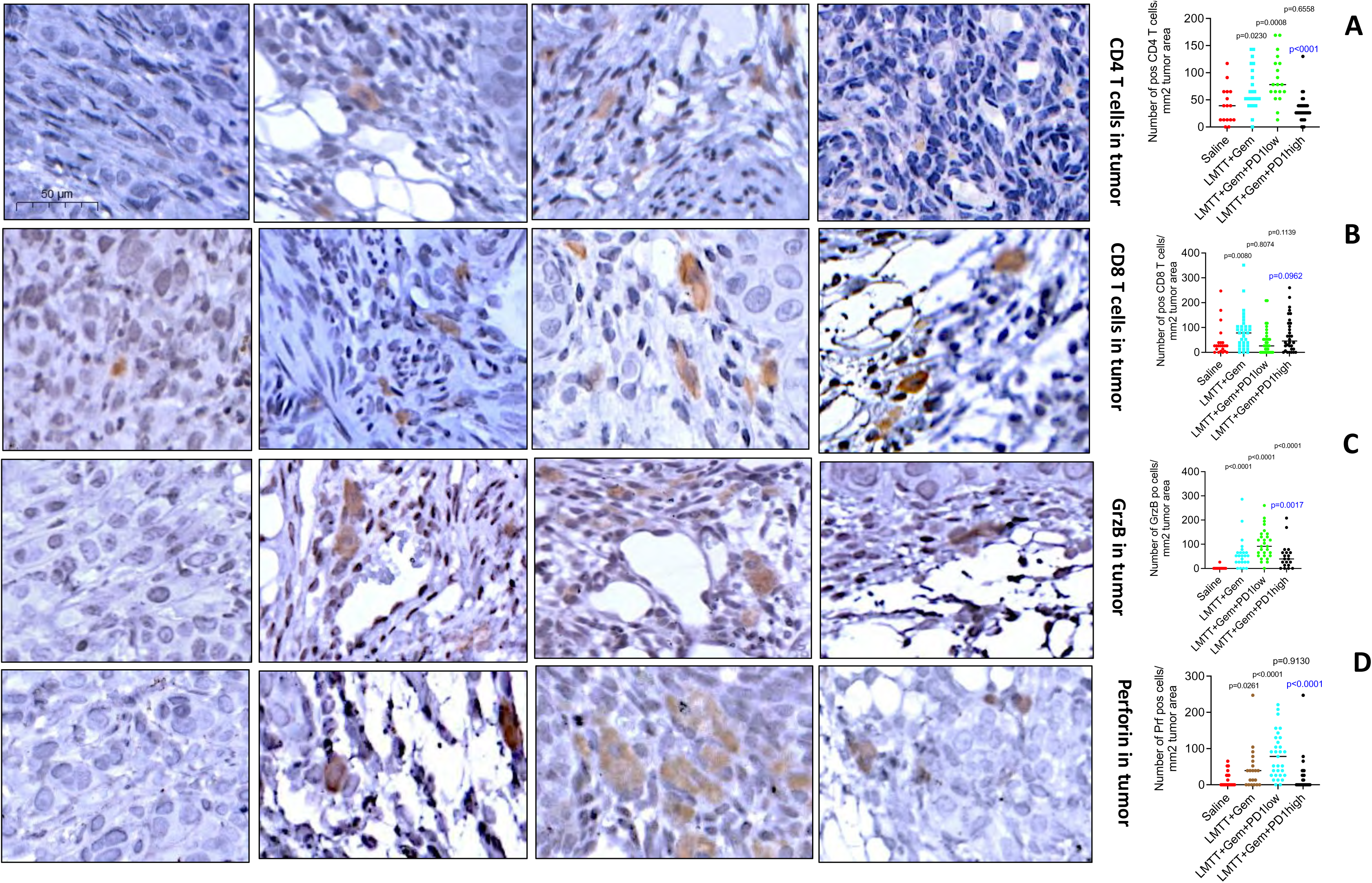
anti-PD1H (PD1H group) inhibited T cell function compared to anti-PD1L (PD1L group), when combined with Listeria-TT+GEM, in the Id8p53-/- Luc model. **(A)** The number of CD4 T cells decreased significantly in the ovarian tumors of PD1H compared to PD1L group using the Id8p53-/- ovarian model. No significant difference was observed in the number of CD8 T cells in the PD1H vs PD1L group. The production of Granzyme B **(C)** and Perforin **(D)** in the Id8p53-/- ovarian tumors significantly decreased in the PD1H compared to the PD1L group. 20 fields per group were counted and the results were averaged. The average number of positive cells were calculated per square millimeter tumor area. n=3 mice per group. Statistical significance was determined by using Mann-Whitney, p<0.05 is significant. All groups in **fig A-D** were compared to the Saline group. Error bars represent SEM. Black p values are all compared to Saline, and blue p value Listeria-TT+GEM+PD1L is compared to Listeria-TT+GEM+PD1H.

### RNAseq analysis of tumors in PD1L vs PD1H group

To obtain mechanistically more insight why high doses of PD1 (PD1H) were less effective than low doses of PD1 (PD1L), when combined with Listeria-TT+GEM, we analyzed gene expression levels and biological pathways in both groups (PD1L vs PD1H) by RNAseq data (see Methods). The analysis revealed that out of 905 genes that were up (+3 to +7-fold difference) or down regulated (−3 to 7-fold difference), we found 18 genes of interest (mostly upregulated) that may reveal mechanisms involved in the weaker effect of the high compared to low doses of anti-PD1 on the ovarian cancer (**figure 9A**). Highest difference in gene expression levels was observed for genes *flg, Ighv-43* and *spink1* in the PD1H group. The upregulation of these genes may contribute to the lower immune infiltration in tumors, more immune suppression, and more cancer stemness. In addition, we performed GSEA analysis of biological pathways and found that the 8 curated pathways (M2) (**figure 9B**) and 7 Gene Ontology pathways (M5:GO) were enriched in the PD1H group (**figure 9C**), separately, while 3 curated pathways were significantly enriched in the PD1L group (**figure 9B**). Most significant were the *Kdm1A*, *DNA repair*, *Immature B lymphocyte, epidermal cell differentiation, and endopeptidase activity* pathways (**figure 9BC**). The enrichment of these pathways suggests involvement of enhanced invasion, less efficient DNA repair systems, less efficient antigen presentation (involving immune responses), and stronger epithelial-mesenchymal transition (EMT) in the PD1H group, resulting in more aggressive ovarian cancer.

**Figure 9:**
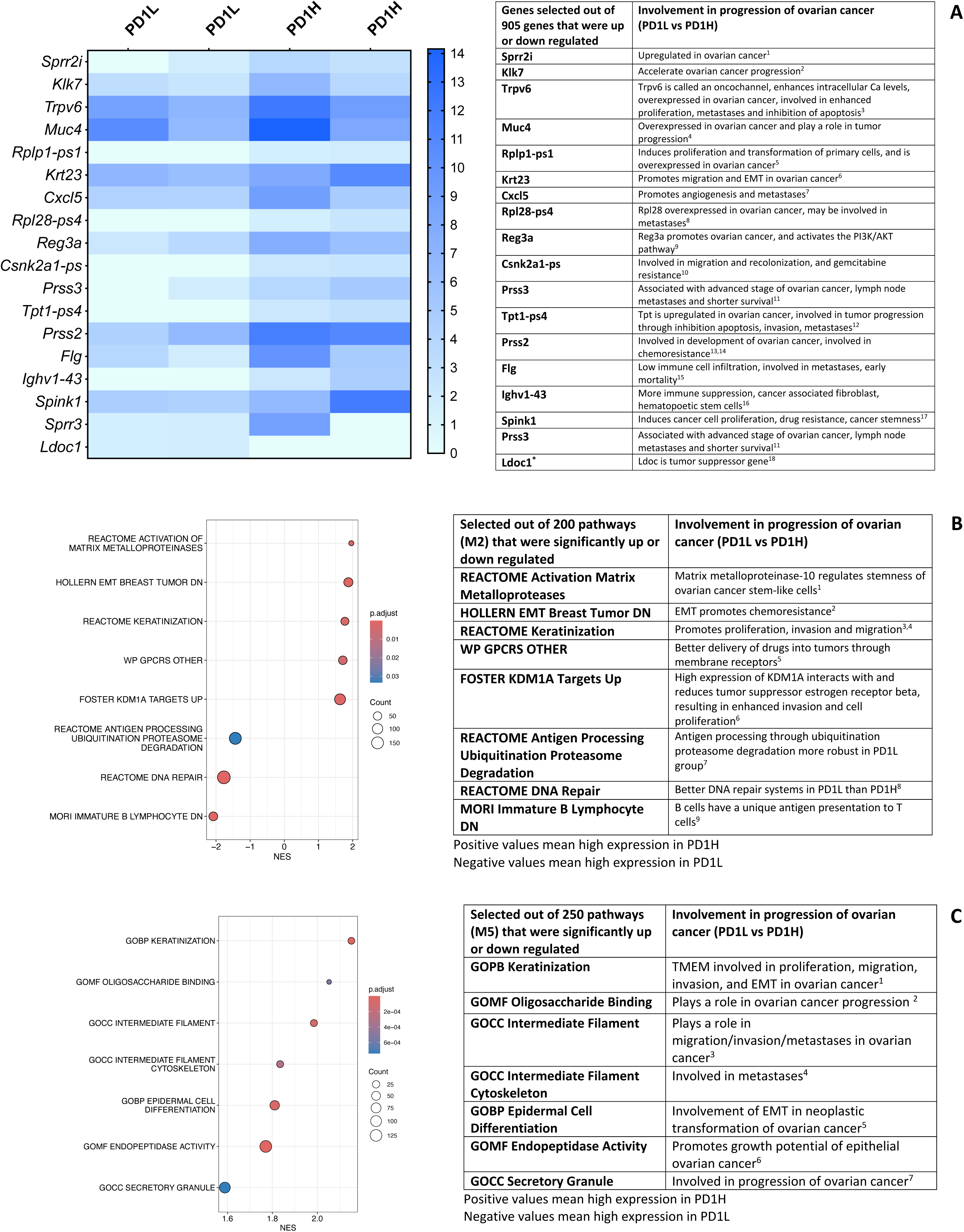
RNAseq analysis of tumors in the PD1L vs PD1H group, combined with Listeria-TT+GEM in Id8p53-/-Luc mice. **(A)** Tumors of ID8p53-/-Luc mice treated with Listeria-TT+GEM combined either with low (PD1L) or high (PD1H) doses of anti-PD1 were analyzed by RNA-seq. The heatmap was generated from the peritoneal tumors by unbiased hierarchical clustering at the following statistical parameters. Statistical parameters for the paired-wise comparison: p < 0.05. The signature contains 18 genes mostly upregulated (Log2 fold difference +3 to +7) or down regulated (Log2 fold difference -3 to -7) in the PD1H compared to the PD1L group. **(B)** and **(C)** The enrichment of biological pathways was analyzed by GSEA; Curated pathways (M2; 8 pathways) were enriched or depleted (NES -2 to +2) in the PD1H compared to the PD1L group, and GO pathways (M2:GO; 7 pathways) were enriched in the PD1H compared to the PD1L group (NES +1.8 to +2.2).

## Discussion

In this study we tested whether Listeria-TT+GEM was effective against ovarian cancer. We generated two Id8p53-/-Luc models. One model predominantly develops metastases in the peritoneal cavity (peritoneal model), and the other model predominantly develops a tumor in the ovaries (ovarian model). The effect of the immunotherapy on the cancer was analyzed by IVIS (since the tumor cells expressed Luciferase), and the survival by a Kaplan-Meier curve. We found that 80-90% of the ovarian tumors expressed TT, which attracted both CD4 and CD8 T cells to the TME, producing perforin and granzyme B. A strong reduction was observed in the primary ovarian tumors compared to the saline group (78%), and inhibition of the growth of peritoneal metastases compared to the saline group (95%), resulting in a significant improvement in survival time compared to all control groups. However, 5% and 22% of the metastases and tumors were left after the therapy, respectively. These tumor cells don’t express TT and will thus not be eliminated by the TT-specific T cells. We hypothesized that delivering more TT to these remaining TT-negative tumor cells by a second treatment cycle of Listeria-TT+GEM, will lead to longer survival. Two treatment cycles doubled (50%) the survival time compared to untreated mice (saline) and increased the survival time by 24% when compared to one treatment cycle. In summary, we were able to strongly improve the survival time, but not to completely eliminate the cancer. These results encourage combining our therapy with other therapies in order to further improve the survival time.

One such therapy is checkpoint inhibitors. Checkpoint inhibitors have been extensively studied in ovarian cancer patients. For instance, PD1 inhibitors (pembrolizumab, dostarlimab) has shown minimal treatment effect in ovarian cancer, partly because ovarian cancer is poorly immunogenic with low neoantigen expression and poor T cell infiltration ^25^. A phase 3 trial using pembrolizumab(anti-PD1) as single-agent in recurrent ovarian cancer showed an overall response rate of only 8%. A phase 2 trial (NCT02718417) testing Avelumab (an anti-PDL1 antibody) plus chemo or avelumab alone showed significant progression-free survival or overall survival versus chemo alone in the platinum resistant/refractory ovarian cancer. Another study examined nivolumab alone (anti-PD1) vs Ipilimumab(anti-CTLA4)+Nivolumab in recurrent ovarian cancer (NCT02498600). Patients treated with Nivolumab alone demonstrated an overall response rate of only 12.2% while the combo showed a response rate of 31.4%. Importantly though, the progression free survival in the nivolumab alone group was 2 months and in the combo group was only 3.9 months which is not clinically significant. These trials demonstrate that checkpoint inhibitors (PD1 or CTLA4) alone were not effective in patients with ovarian cancer. As mentioned above, such failure is partly because ovarian cancer is poorly immunogenic. Also, in our mouse models we found that anti-PD1 alone was also poorly effective. We hypothesized that if we deliver our own neoantigen surrogate (TT) to the TME, it may improve the effect of checkpoint inhibitors.

We first tested two different doses of anti-PD1 antibodies. It appeared that the low dose (50 μg/dose) was more effective than the high dose (130 μg/dose) anti-PD1. We then repeated the experiments with low doses of anti-PD1 and found that Listeria-TT+GEM+anti-PD1 was significantly more effective than anti-PD1 or Listeria-TT+GEM alone, as shown by IVIS and ROI counts quantified, and supported by survival data.

From a clinical perspective, the high vs low doses of anti-PD1 is an important issue because of the immune related adverse events (irAE), often described for PD1 inhibitors. This includes autoimmune-like/inflammatory side effects, causing collateral damage to normal organ systems and tissues such as skin, GI, hepatic, pulmonary, mucocutaneous, and endocrine systems)^26^. Therefore, we investigated why the low doses were more effective than the high doses, i.e. we evaluated the effect of high and low doses of anti-PD1 on the Listeria viability (growth), immune function, and on potential toxicities on organs (H&E staining). Very mild toxicities were observed in the tissues (liver, spleen, lungs, kidneys and GI) with both high and low doses of anti-PD1 antibodies. Also, no effect of the high or low doses of anti-PD1 was observed on the viability of Listeria. However, we found an inverted dose-response effect of anti-PD1 antibodies on spleen cells of TT-vaccinated mice, infected with Listeria-TT in the presence of various concentration of anti-PD1 antibodies. The higher the anti-PD1 dose, the more inhibition of the production of IFNγ in the spleen was observed. This correlated with the *in vivo* results, i.e. higher doses of anti-PD1 were less effective against the tumors and metastases *in vivo*. We analyzed this in more detail by IHC and found that high doses of anti-PD1 attracted significantly less CD4 T cells to the TME compared to low anti-PD1, producing significantly less perforin and granzyme B, when combined with Listeria-TT+GEM, suggesting that high doses of anti-PD1 inhibited T cell function and low doses of anti-PD1 stimulated T cell function.

To obtain more mechanistic insight in the efficacy of the PD1L vs PD1H treatments, we analyzed gene expression levels and biologic pathways enrichment in the ovarian tumors of both groups. The DEG analysis showed upregulation of genes involved in less immune infiltration and more immune suppression, resulting in more aggressive cancer in the PD1H group, while the GSEA on biological pathways pointed towards less efficient antigen processing and presentation, and less efficient DNA repair systems, as well as a stronger EMT, resulting in more aggressive tumors in the PD1H group. Although immune responses were definitely less potent in the PD1H compared to the PD1low group in our mice, pathways involved in migration, invasion, metastases seem more dominant.

A question that arises from these experiments is whether we use the optimal dose of PD1 in patients. In patients, dose escalation studies with checkpoint inhibitors have been performed. However, this only involved toxicity but not efficacy. Of note, the higher doses of PD1 in our study did not lead to higher toxicity either. Thus, in patients, we don’t know if we use the optimal efficacy dose. The clinical importance of this issue may justify further analysis.

Others have studied checkpoint inhibitors in combination with CAR T cells in mice. In one study, human Muc16-specific CAR T cells combined with anti-PD-L1 in NSG mice bearing OVCAR3 tumors expressing Muc16, PD-1 and Luc^27^. The tumor cells (OVCAR3) were transduced with muc16 tumor antigens and PD1 and Luc *in vitro*. Results were highly promising as the survival time was significantly improved in the treatment group as compared to the control mice ^27^. In another study they combined CAR T cells against mesothelin with anti-PD1 ^28^. In this study, SKOV3 and OVCAR4 human tumor cells were transduced *in vitro* with mesothelin antigen-GFP-Luc, which is an endogenous tumor antigen. Their CAR T cells, when combined with anti-PD-L1, were able to control the growth of SKOV3 and OVCAR4 tumors and significantly improved the survival time ^28^. These results support our data because overexpression of antigens by the tumor increased the effectiveness of anti-PD1 or anti-PD-L1 therapies. However, while these studies are highly promising, it needs to be noted that their antigens were delivered into the tumor cells *in vitro* through transduction, which is impossible to do with patients. This is completely different from our approach, where we deliver TT antigen *in vivo* in the tumors through Listeria infection. Moreover, in contrast to relative poorly immunogenic tumor antigens, TT antigens are functioning as a neoantigen surrogate (highly immunogenic) and as a recall antigen (to which the immune system has been exposed to earlier in life through childhood vaccinations).

## Conclusions

In summary, the results of this study suggest that patients with HGSC may benefit from our approach. Currently, we are preparing for a Phase 1 clinical trial of Listeria-TT+GEM in patients with advanced PDAC. If successful, we will follow up with HGSC patients.

## Methods

### Animals

C57BL/6 mice (females only) of 6-8 weeks were obtained from Charles River. These mice were used to generate mice with orthotopic Id8p53-/- Luc tumors in the ovaries (orthotopic model) and mice with metastases in the peritoneal cavity (peritoneal model).

### Cell lines

The Id8p53-/- Luc cell line is a high grade serous ovarian carcinoma (HGSC) model when injected into the peritoneal cavity. This cell line was developed and kindly provided by Dr. Mc Neish ^16^, Centre for Molecular Oncology, Barts Cancer Institute, Queen Mary University of London, and Dr. Janet Sawicki, Sidney Kimmel Cancer Center, Thomas Jefferson University, Philadelphia, Pennsylvania ^17^. The HEY tumor cell line was derived from human HGSC ^29^ and was kindly provided by Dr. Susan Horwitz, Albert Einstein College of Medicine, Bronx. This cell line was used to determine infection and destruction by Listeria-TT *in vitro*.

### Mouse models

In this study two mouse models were generated, i.e. a peritoneal model and an ovarian model. One model consisted of injection of the Id8p53-/-Luc cells in the peritoneal cavity, which resulted in metastases in the peritoneal cavity but no tumors in the ovaries or uterus, while the other model consisted of injection of Id8p53-/-Luc cells in the ovaries resulting in primary tumor in the ovaries. In both cases, tumor growth was measured by In Vivo analysis of Small animals (IVIS) as we described earlier ^30^.

### Peritoneal model

Metastases were generated in immune competent C57BL/6 mice by injection of 1-2x10^6^ Id8p53-/-Luc cells/200uL DPBS into the peritoneal cavity. 10-14 days after the injection, when metastases were developed, the immunotherapeutic treatments were started. Metastases were visible in the portal vein area of the liver, the diaphragm, along the GI and in the pancreas (no tumors were developed in the ovaries).

### Ovarian model

Orthotopic tumors were generated in the ovaries by injection of 2x10^6^ Id8p53-/- tumor cells in the ovaries of immune competent C57BL/6 mice. Briefly, mice were anesthetized with ketamine (Mylan Institutional LLC/xylazine (Akorn Animal Health)(respectively, 100 mg and 10mg/kg, ip)^9^, the hair was removed at the location above the hindleg, and the skin was sterilized with betadine, followed by 70% alcohol. The animal was covered with gauze sponge surrounding the incision site. A 1 cm incision was made in the abdominal skin above the right hindleg to allow visualization of the ovaries and uterus. Carefully, the uterus was uplifted by forceps and 2x10^6^ Id8p53-/-Luc tumor cells mixed 1:1 with Matrigel (VitroGel Hydrogel Matrix, MidSci, Scranton, PA; cat# VHM01S) in 50 μl volume were injected into the bursa of the ovary. The ovary was then replaced within the abdominal cavity, and both skin and muscle layers were closed with sutures. Following recovery of the surgery, mice were monitored and weighed daily. The ovarian tumor was visible by IVIS within 10-14 days. The tumor growth was measured by IVIS before (day 0 of treatment), during (day 10), and after treatment (day 20).

### Listeria and Listeria-TT

In this study, an attenuated *Listeria monocytogenes* (*Listeria*) was used as the vehicle for delivery of TT856-1313 ^31^ to the TME and inside tumor cells. The *Listeria* plasmid pGG-34, expressing the non-cytolytic truncated form of Listeriolysin O (LLO) under control of the hemolysin promoter *(Phly)* and the positive regulatory factor A (*prfA*), with mutations to further reduce pathogenicity, have been described elsewhere ^32^. *Listeria*-TT_856-1313_ was developed in our laboratory as described previously ^9^. The TT_856-1313_ fragment contains both mouse and human immunodominant T cell epitopes ^33,34^.

### Biodistribution of *Listeria*-TT

C57BL/6 mice were injected intraperitoneally with 2x10^6^ Id8p53-/- mouse ovarian tumor cells as described above. To visualize big metastases in the peritoneal cavity we injected with a single large dose 10^7^ CFU of *Listeria*-TT ip, 4 weeks after tumor cell injection. At this time point the mice developed visible metastases by eye and ascites. The ascites was drained from live mice before injection of the Listeria-TT. Mice were euthanized at various time points after the injection (as indicated in the figure), and metastases and normal tissues were dissected, weighed, and analyzed for CFU of *Listeria*-TT, as described previously ^15^.

### Infection of tumor cells *in vitro*

The *in vitro* infectivity of Id8p53-/- (mouse) or Hey tumor (human) ovarian cancer cells was assessed as described previously ^14^. Briefly, 10^6^ tumor cells/ml were infected with 10^7^ CFU of *Listeria* or *Listeria* -TT for 1hr at 37°C in culture medium, then incubated with gentamicin (50 μg/ml) for 1 hr to kill extracellular *Listeria*. Finally, cells were washed with phosphate-buffered saline (PBS), lysed in water, and serial dilutions were plated onto LB agar to quantify the infection rate the next day.

### Infection of tumor cells *in vitro* in the presence of various doses of anti-PD1 antibodies

We also tested the effect of high and low doses of anti-PD1 antibodies on the infection rate of Id8p53-/- cells by Listeria-TT. Briefly, the infection was performed as described above but now in microtiter wells (200 μL) in the presence of serial dilutions of high or low doses of anti-PD1 (low dose 50, 5, and 0.5 μg/well and high dose 130, 13, and 1.3 μg/well). After 1hr culture, Gentamicin was added (50 μg/mL) for 1 hr, and subsequently the bacteria were isolated as described above.

### Evaluation of cell death

As described previously ^14^, tumor cell killing by *Listeria* or *Listeria*-TT was determined *in vitro* as follows. Id8p53-/- or Hey tumor cells (3x10^3^) plated in 96 well plates were infected with 10^7^ CFU/well of Listeria or Listeria-TT for 2hrs at 37°C, then gentamicin (50 μg/ml) was added. Live and dead cells were counted the next day using Trypan blue staining.

### Protocol for Listeria-TT+GEM treatment of Id8p53-/-Luc mice (efficacy and survival studies)

A detailed rationale for this treatment protocol and schematic view of all treatments are shown in **figure S3A.** To generate memory T cells to TT, C57BL/6 mice were immunized twice intramuscularly with the TTvacc on days 0 and 7 (the same used for childhood vaccinations, 0.3 μg/50 μL). Subsequently, Id8p53-/-Luc tumor cells were injected intraperitoneally (1-2x10^6^/200 μL, as indicated in the legends) or orthotopically (2x10^6^/50 uL) in the bursa of the ovaries on day 10-14 (as indicated in the legends), and after we confirmed the presence of metastases or tumor by IVIS, we started the treatments. High doses of *Listeria*-TT (10^7^ CFU, 3 doses total) were injected ip on days 28, 34 and 43 of a complete treatment cycle to deliver TT to the TME. Low doses of GEM (Fresenius kabi oncology Ltd) were injected on days 29, 32, 35, 38, 41, and 44 (1.2 mg/mouse, every 3 days) (six doses in total) to stimulate the TT specific T cells. Concomitantly, low doses of *Listeria*-TT were administered on days 30, 31, 33, 36, 37, 39, 40, and 42 (10^4^ CFU, 8 doses in total) to reduce immune suppression. Growth of tumors and metastases were analyzed by IVIS before (day 27=day 0 of treatment), during (day 37=day 10 of treatment) and after treatment (day 47=day 20 of treatment). Subsequently, all mice were monitored in a survival study (Kaplan-Meier curve). At the end of treatments mice were monitored without any further treatment until they succumbed spontaneously or were terminated upon appearance of severe pre-morbid symptoms requiring euthanasia as specified by our approved animal use protocol. Results were analyzed by Mantel-Cox test.

### Protocol for Listeria-TT+GEM+anti-PD1 treatment of Id8p53-/-Luc mice (efficacy and survival studies)

In the Id8p53-/-Luc model, mice were immunized twice with Tetanus vaccine (TTvac) to generate memory T cells (days 0 and 7). Subsequently, Id8p53-/-Luc tumor cells were injected intraperitoneally or orthotopically in the ovaries (day 14) and metastases and tumors allowed to develop to an advanced stage detectable by IVIS and H&E staining. Three high doses of Listeria-TT were administered on days 28, 34, an 41, combined with four low doses of Gemcitabine on days 29, 32, 37, and 41, and four low dose of Listeria-TT on days 30, 33, 38, and 42, followed by anti-PD1 antibodies on days 31, 35, 39 and 43. Anti-PD1 antibodies were used to induce more durable effects on T cells. Tumors and metastases were monitored by In Vivo Imaging System (IVIS) *in vivo* before (day 27), during (day 37) and after the therapy (day 47). At the end of treatments, we continued monitoring until death. LM-TT=Listeria-TT. A schematic view of the treatment protocol has been shown in **figure S3B.**

### In Vivo Imaging System (IVIS)

C57BL/6 mice were injected with 2x10^6^ Id8p53-/-Luc cells (in 50 μl DMEM) into the peritoneal cavity or ovaries as described above. Ip injection results in metastases in the peritoneal cavity, while intra-ovary injections result in a primary tumor in the ovaries. Metastases and primary tumors were visible by IVIS by 10-14 days after injection. The luciferase activity was measured using an intensified charged-coupled device video camera of the In Vivo Imaging System (IVIS 100; Xenogen). The Id8p53-/-Luc tumor-bearing mice were injected ip with 200 μl of D-luciferin sodium salt (Synchem OHG; catalogue number BC218) dissolved in phosphate-buffered saline (100 mg/kg of body weight). Luciferine distributed systemically for 3 min while the animals were anesthetized (mixture of isoflurane and oxygen). Anesthetized animals were placed in the camera chamber, and a bioluminescent signal was acquired for 5 min. Bioluminescence measurements produced by the IVIS 100 system are expressed as a pseudocolor on a gray background, with red denoting the highest intensity and blue the lowest. To quantify the luminescence, we outlined a region of interest (ROI) and analyzed it by use of the Living Image (version 2.11;Xenogen) and Igor Pro (version 4.02A; WaveMetrics) software.

### Immunohistochemistry (IHC) and H&E staining

Tumors were dissected from the ovaries and immediately fixed with buffered formalin, and the tissue was embedded in paraffin. Sections (5 µm) were sliced and placed on slides, then deparaffinized at 60°C for 1 hr, followed by xylene, an ethanol gradient (100-70%), water, and PBS. Slides were then incubated for 30 min in 3% hydrogen peroxide followed by boiling in citrate buffer for 20 min. Once the slides were cooled, washed, and blocked with 5% goat serum, the sections were incubated with primary antibodies such as anti-CD4 (1:100 dilution), anti-CD8α (1:400 dilution), anti-Perforin (1:300 dilution), anti-Granzyme (1:200 dilution), anti-PD-L1 (dilution 1:100), anti-TT (1:50 dilution), followed by incubation with secondary antibody (mouse anti-goat IgG-HRP), and SignalStain® Boost IHC Detection Reagent (Cell Signaling Technology, cat# 8114S). Subsequently, the slides were incubated with 3,3’-diaminobenzidine (DAB) (Vector Laboratories, cat# SK-4100), counterstained with hematoxylin, dehydrated through an ethanol gradient (70-100%) and xylene, and mounted with Permount. The slides were scanned with a 3D Histech P250 High Capacity Slide Scanner to acquire images and quantification data. Secondary antibodies without primary antibodies were used as negative control. In addition, sections were stained with H&E as we described previously. Briefly, tumors were fixed in 10% formaldehyde for 48 h, and then kept in 70% ethanol until use. Sections of 1 mm thick were stained with hematoxylin/eosin and analyzed to confirm the presence of tumors and metastases.

### ELISPOT in the presence of various doses of anti-PD1 antibodies

Spleen cells were isolated from C57BL/6 mice that received TT vaccinations twice, one week apart for ELISPOT (BD Biosciences, cat# 551083) analysis, as described previously ^35^. To determine immune responses to TT, 2x10^5^ spleen cells were incubated with Listeria-TT (1:5). To measure the effect of anti-PD1 antibodies on immune function of spleen cells (production of IFNγ), anti-PD1 (Biolegend cat#135247) was added to the spleen cells at various doses (low dose 50, 5, and 0.5 μg/well and high dose 130, 13, and 1.3 μg/well). The frequency of IFNγ-producing spleen cells was measured 48 hrs later using an ELISPOT reader (CTL Immunospot S4 analyzer).

### RNAseq

#### RNA Preparation and Extraction

5 tumor sections, each 5 um thick per mouse, were pooled and sent to Zymo Research. By using the Quick-RNA FFPE Miniprep (R1008), total RNA was purified and treated with DNase I. Nanodrop and Agilent’s RNA ScreenTape Assay on TapeStation were used to performed quality control on two mice in each group (Group I: Listeria-TT+GEM+anti-PD1L and Group II: Listeria-TT+GEM+anti-PD1H). 240 ng of Total RNA were used to construct libraries.

#### RNA-Seq Library Preparation

Libraries were prepared using the Zymo-Seq RiboFree Total RNA Library Prep Kit (Cat # R3000) according to the manufacturer’s instruction manual v1.3.0. Briefly, RNA was reverse-transcribed into cDNA, which was followed by ribosomal RNA depletion. After that partial P7 adapter sequence was ligated at 3’ end of cDNAs, followed by second strand synthesis and partial P5 adapter ligation to 5’ end of the double stranded DNAs. Lastly, libraries were amplified to incorporate full length adapters under the following conditions: initial denaturation at 95°C for 10 min; 10 - 16 cycles of denaturation at 95°C for 30 sec, annealing at 60°C for 30 sec, and extension at 72°C for 60 sec; and final extension at 72°C for 7 min. Successful library construction was confirmed with Agilent’s D1000 ScreenTape Assay on TapeStation. RNA-Seq libraries were sequenced on an Illumina NovaSeq X Plus to a sequencing depth of at least 30 million read pairs (150 bp paired-end sequencing) per sample.

#### RNA-Seq Data Bioinformatics Analysis

Low-quality bases and poly-G at the read tail are trimmed by fastp (v0.23.4)^36^. Only reads with length larger than 50 bp were kept for alignment. The trimmed reads were aligned to the mouse reference genome (build GRCm38.102) using STAR (v2.7.11b)^37^. The read counts were quantified with featureCounts (v2.0.6)^38^. The downstream analysis was performed on tumors from Id8p53-/-Luc mice treated with *Listeria*-TT+GEM+anti-PD1L or anti-PD1H. The differentially expressed genes were identified by DESeq2 (v1.42.0)^39^. The gene set enrichment analysis was performed through clusterProfiler (v4.10.0)^40^. We obtained curated gene sets (M2) and gene ontology gene sets (M5: GO) from MsigDB^41^. Specially, M2 include gene sets from CGP (chemical and genetic pertubations), BioCarta, Reactome, and WikiPathways.

### Statistics

To statistically compare the effects of *Listeria*-TT+GEM with or without anti-PD1 antibodies on the growth of metastases and tumors or on immune responses in the mouse models, a non-parametric Mann-Whitney was applied using Prism. The type of statistical test was determined by prism. \**p*<0.05, **<0.01, ***<0.001, ****<0.0001. Values of *p*<0.05 were considered statistically significant. To statistically compare the effects of *Listeria*-TT+GEM on IHC and efficacy Mann-Whitney (two-tailed) test was used. To statistically compare the effects of *Listeria*-TT+GEM with or without anti-PD1 antibodies in a survival study a Kaplan-Meier curve was generated using Mantel-Cox . \**p*<0.05 is significant.

## ETHICS APPROVAL AND CONSENT TO PARTICIPATE

Animal protocol approval number 00001313 according the guidelines of the Institutional Animal Care and Use Committee (IACUC) at Albert Einstein College of Medicine (expiration date 11/11/2026). Human tissues or blood or clinical trials: Not Applicable.

## CONSENT FOR PUBLICATION

All authors have agreed to publish this manuscript.

## DATA AVAILABILITY

All the data are present in the manuscript or in the Supplementary Figures.

## Competing interests

CG. is the inventor of the Listeria-Recall Antigen Technology, which was developed in her laboratory, and is described in a patent application 11213577 (granted in the United States, China and Japan). The patent is licensed to Loki Therapeutics. CG. is a stockholder of Loki Therapeutics and is an employee of Albert Einstein College of Medicine. All other authors declare that they have no competing interests.

## AKNOWLEDGEMENTS

This work was supported by a NCI R21 grant 1R21CA292190-01, and Loki Therapeutics. Drs. Kuo, Isani, and Nevadunsky, Division of Gynecologic Oncology Montefiore/Einstein provided the clinicians (surgeons of ovarian cancer) performing the experiments in Dr. Gravekamp’s lab. The NCI cancer center grant P30CA013330 supported multiple core facilities at Einstein used for this manuscript (Flow Cytometry Core, Histotechnology and Comparative Core Facility, In Vivo Imaging System, Animal housing, Analytical Imaging Core, and the biostatics Shared Core), and the Shared Instruments Grant (Flow cytometry 1S10OD026833-01, IVIS 1S10OD034192-01A1, and 3DHISTECH P250 slide scanner 1S10OD026852-01A1). This study was also supported by Katie Chudnovsky (private donation).

## AUTHOR’S CONTRIBUTIONS

LRS, LG, OK, SA, EL, EB, AQ, SS and CG performed experiments, analyzed the data, and prepared the manuscript. KL, DYSK SI, SS, and NN contributed intellectually to the manuscript, while CG conceptualized and supervised the project, revised the manuscript, and provided the resource for this project.

## A SUPPLEMENTARY INFORMATION

**Figure S1:**
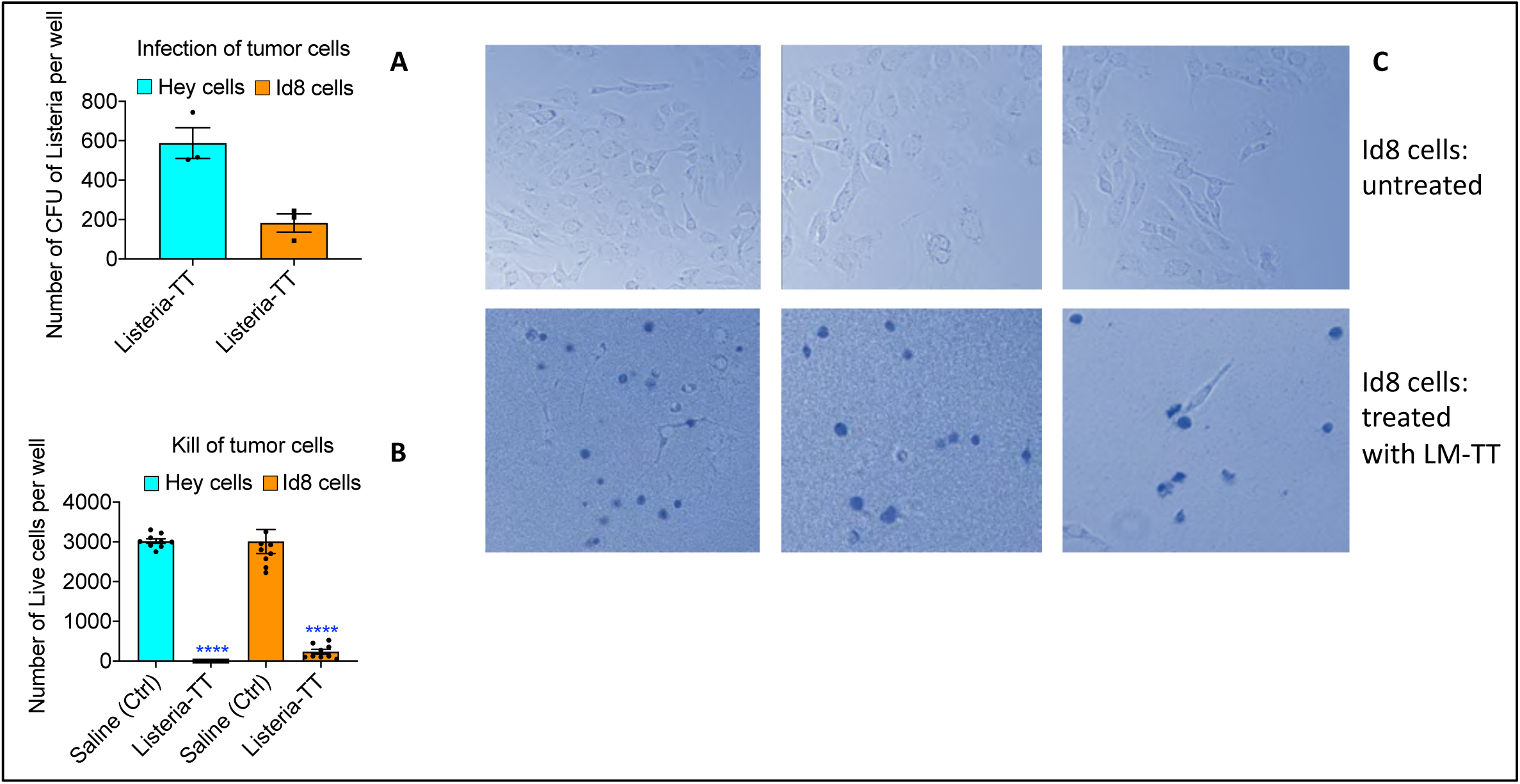
Listeria-TT infecting and killing mouse and human ovarian cancer cells. A, Listeria-TT, infected both human chemoresistant Hey and mouse ID8p53-/- ovarian cancer cells *in vitro* with high efficiency after one hour of infection. Experiments were performed in triplicates, and repeated three times. The results presented in this figure are the average of three infection experiments. The error bars represent the SEM. **B,** We also tested the killing of Hey and Id8p53-/- tumor cells by Listeria-TT and found that both were efficiently killed by Listeria. In this experiment, tumor cells were incubated with the *Listeria* bacteria for 2 hours and then treated with gentamycin. **C,** Next day, dead and alive tumor cells were determined with trypan blue. From each sample, nine fields were blinded analyzed by two different investigators. Experiments were performed in triplicates, and repeated three times. The results presented in this figure are the average of three experiments. The error bars represent SEM.

**Figure S2:**
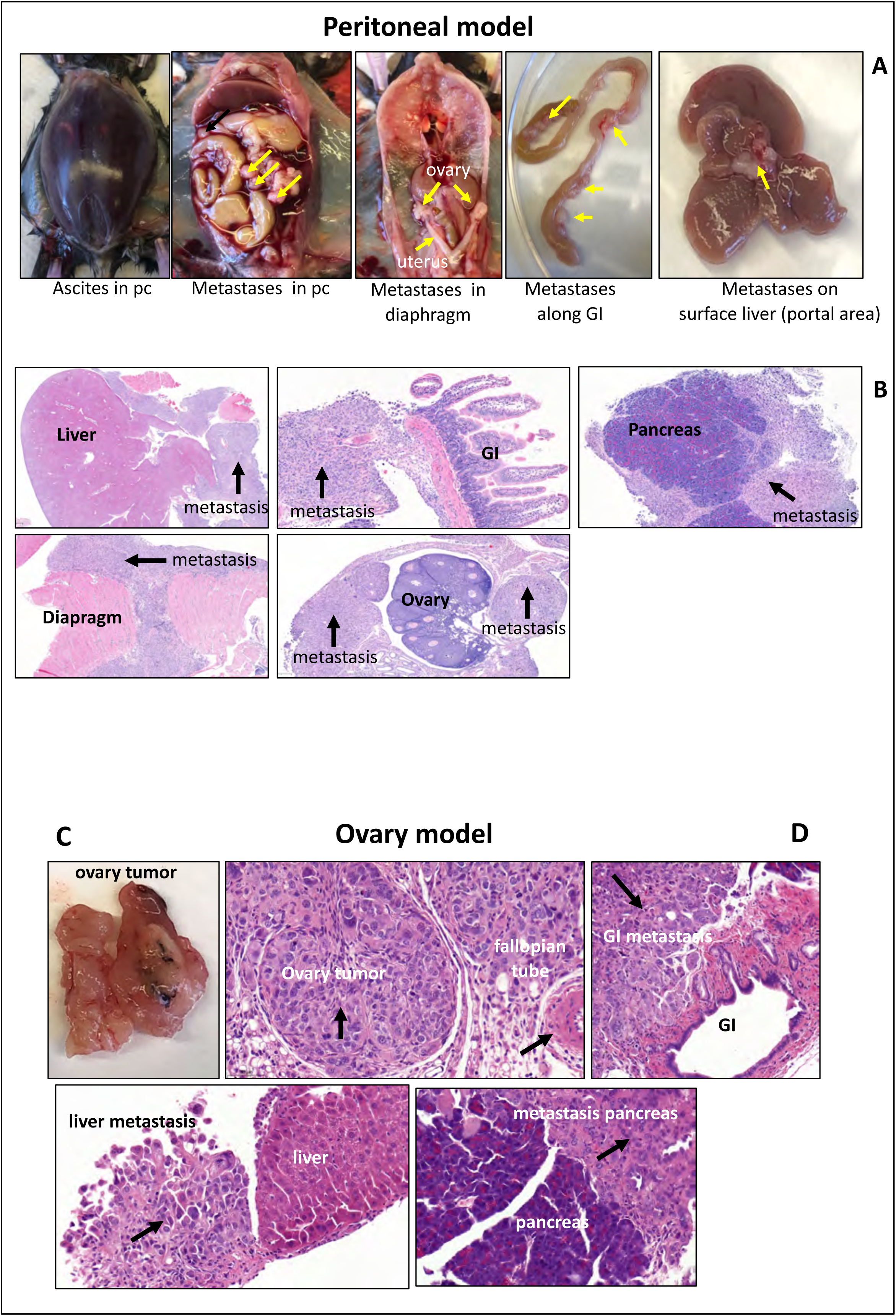
Characteristics of the peritoneal and ovary Id8p53-/- models. Two ovarian cancer models were tested, i.e. the peritoneal and the ovary model. Both models were analyzed by pathological examination to confirm the cancer. In the peritoneal model, tumor cells were injected intraperitoneally resulting in metastases predominantly in the peritoneal cavity along the GI, in the diaphragm, and in the area of the portal liver visible by the naked eye **(A)** and by pathological analysis (H&E staining) **(B).** No tumor cells were detected in the ovaries. In the ovary model, tumor cells were injected into the ovaries, resulting in tumors predominantly in the ovaries visible by the naked eye **(C)** and by pathological analysis (H&E staining) **(D).** Metastases were only found 4-5 weeks later.

**Figure S3A:**
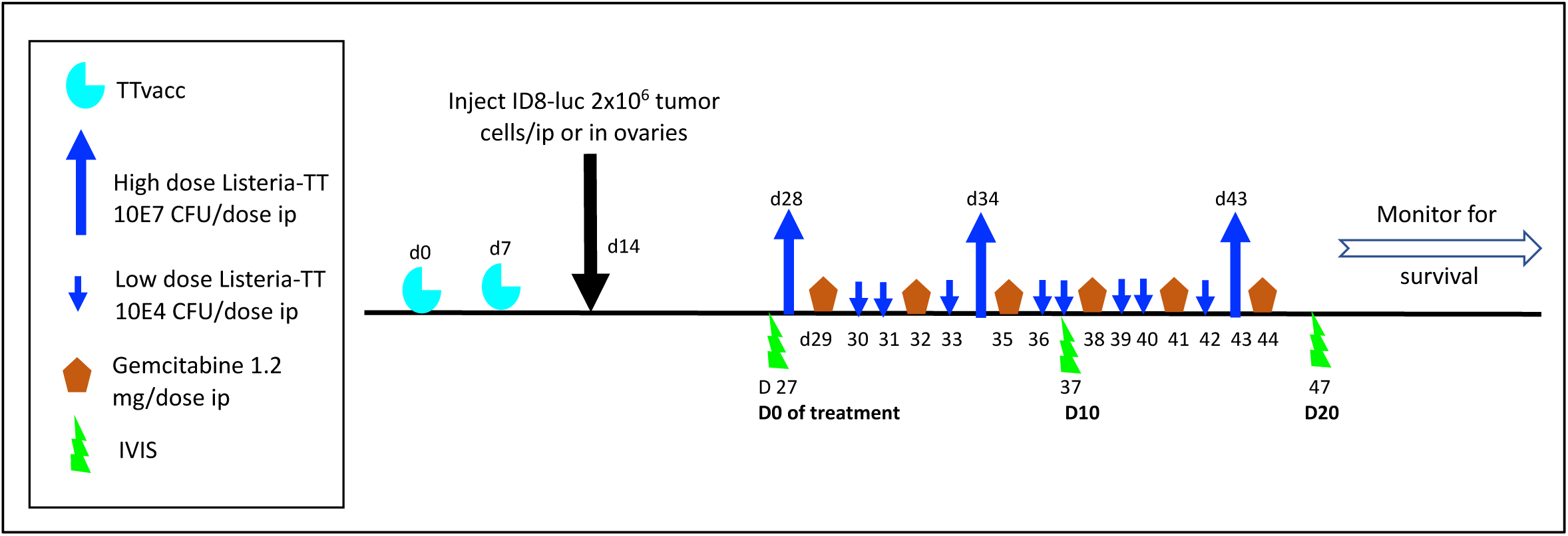
Treatment protocol Listeria-TT+GEM. C57BL/6 mice were immunized twice (days 0 and 7) with the Tetanus Toxoid vaccine (TTvacc) to generate memory T cells to TT. Subsequently, Id8p53-/-Luc tumor cells were injected intraperitoneally or orthotopically in the ovaries (day 14) and metastases and tumors allowed to develop to an advanced stage detectable by IVIS and H&E staining (day 28). After an initial high ip dose of *Listeria*-TT (10^7^ Colony-Forming Units (CFU)(day 28), Listeria-TT accumulated in the TME (within the first 4 hrs after injection). Low doses of GEM (1.2 mg/dose; 6 doses total on days 29, 32, 35, 38, 41, and 44) were then given every 3 days for 14 days in order to eliminate MDSC and TAM. At this point, MDSC are no longer required to bring *Listeria*-TT to the TME. Instead, reducing MDSC and TAM populations by GEM serves to improve T cell responses. Two additional high doses of Listeria-TT were administered to deliver more TT at the tumor site on days 34 and 43. Alternately, daily low doses of *Listeria*-TT (10^4^ CFU; 10 doses total on days 30, 31, 36, 37, 39, 40, 42) to reactivate TT-specific memory T cells, improved by GEM. Tumors and metastases were monitored by intravital imaging of small animals in vivo (IVIS) before (day 27), during (day 37) and after the therapy (Day 47). At the end of treatments, we continued monitoring until death. LM-TT = *Listeria*-TT.

**Figure S3B:**
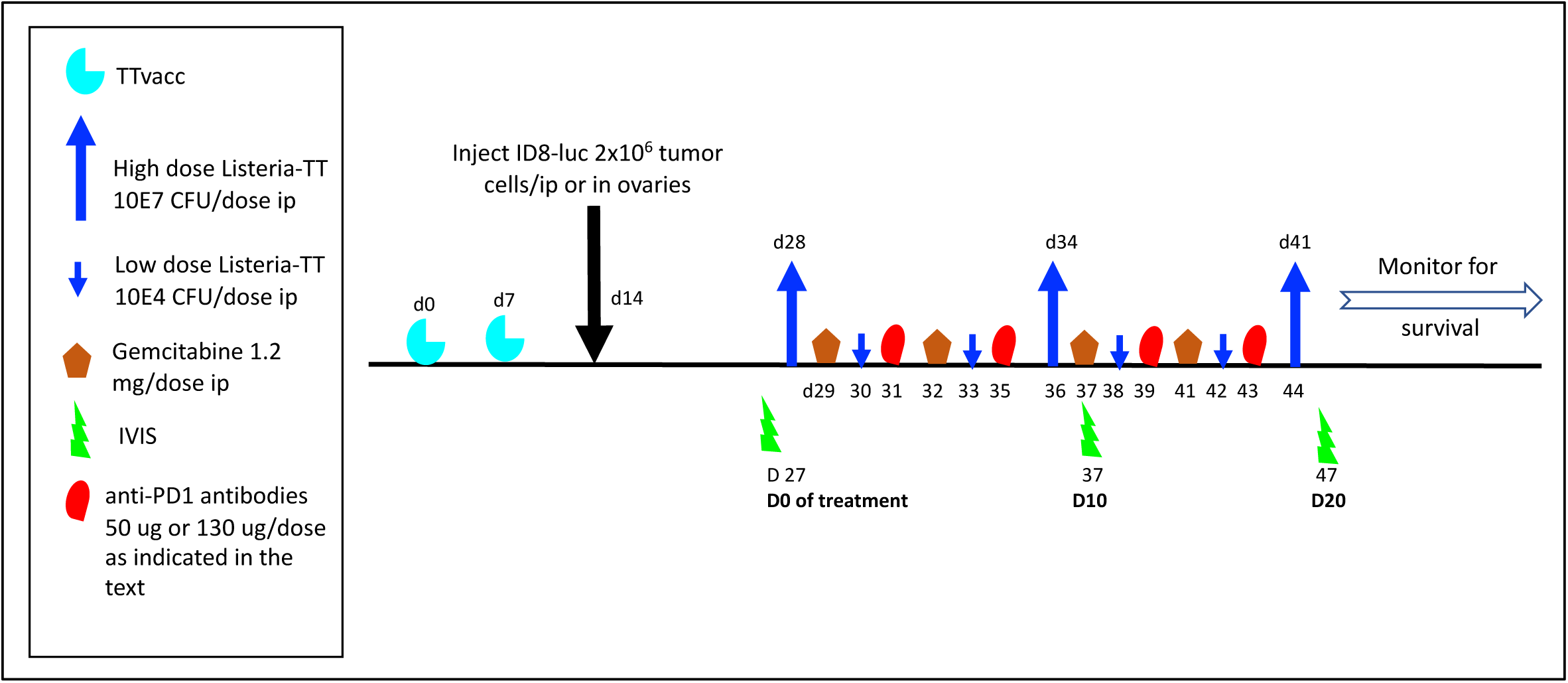
Treatment protocol Listeria-TT+GEM+anti-PD1. In the Id8p53-/-Luc peritoneal model, mice were immunized twice with Tetanus vaccine (TTvac) to generate memory T cells (days 0 and 7). Subsequently, Id8p53-/-Luc tumor cells were injected intraperitoneally (day 14) and metastases and tumors allowed to develop to an advanced stage detectable by IVIS and H&E staining. Three high doses of Listeria-TT were administered on days 28, 34, an 41, combined with four low doses of Gemcitabine on days 29, 32, 37, and 41, and four low dose of Listeria-TT on days 30, 33, 38, and 42, followed by anti-PD1 antibodies on days 31, 35, 39 and 43. High (130 μg/dose) and low doses (50 μg/dose) of anti-PD1 were tested as indicated in the text. Tumors and metastases were monitored by intravital imaging of small animals in vivo (IVIS) before (day 27), during (day 37) and after the therapy (Day 47). At the end of treatments, we continued monitoring until death. LM-TT=Listeria-TT

**Figure S4AB:**
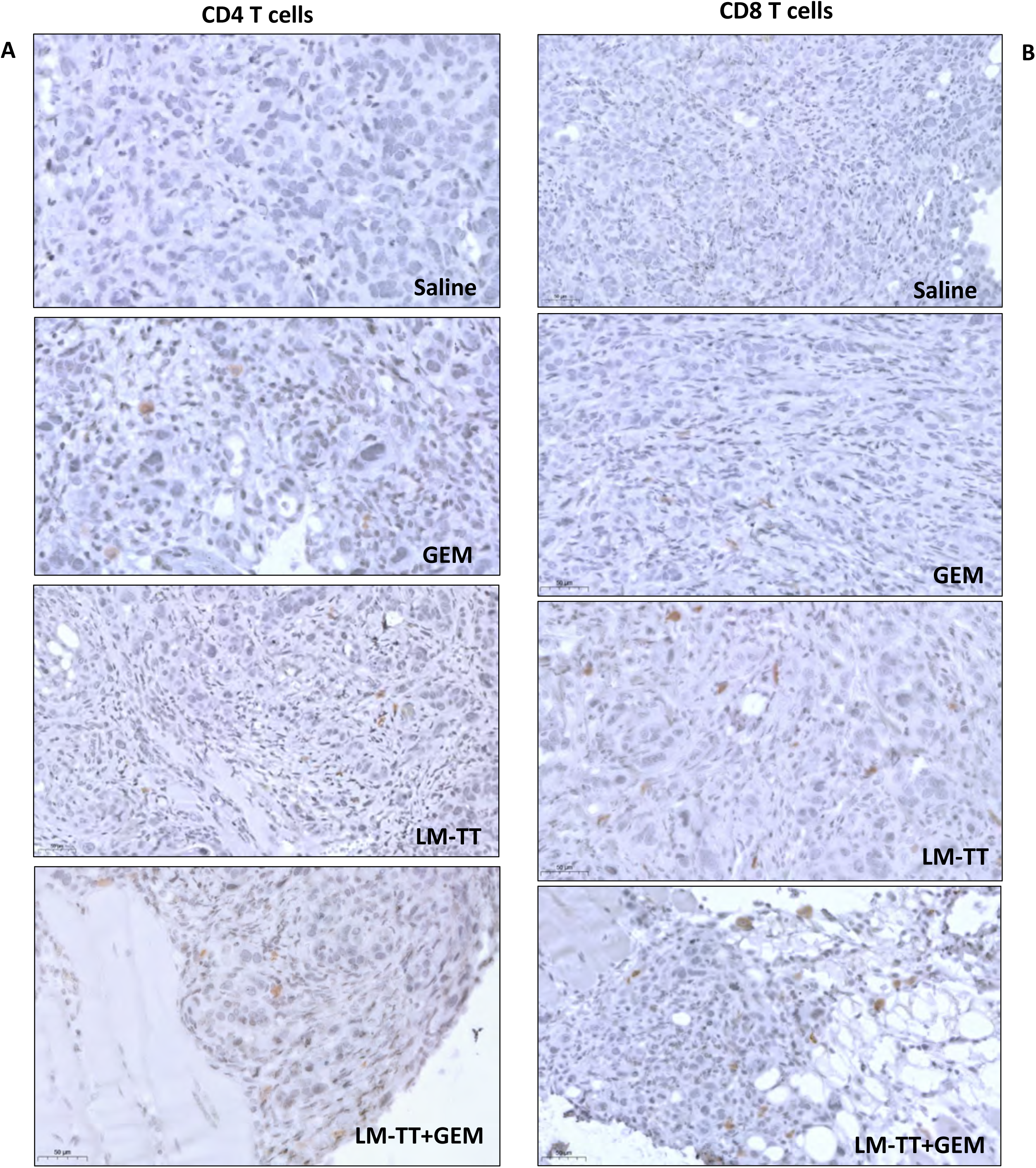
CD4 T cells and CD8 T cells in ovary tumors. Id8p53-/-Luc mice (ovary model) received treatments with Listeria-TT+GEM, or control treatments, as outlined in Figure S1A. **AB,** Two days after the last treatment, tumor tissues were analyzed by immunohistochemistry (IHC) for the presence of CD4 and CD8 T cells. LM-TT = Listeria-TT.

**Figure S4CD:**
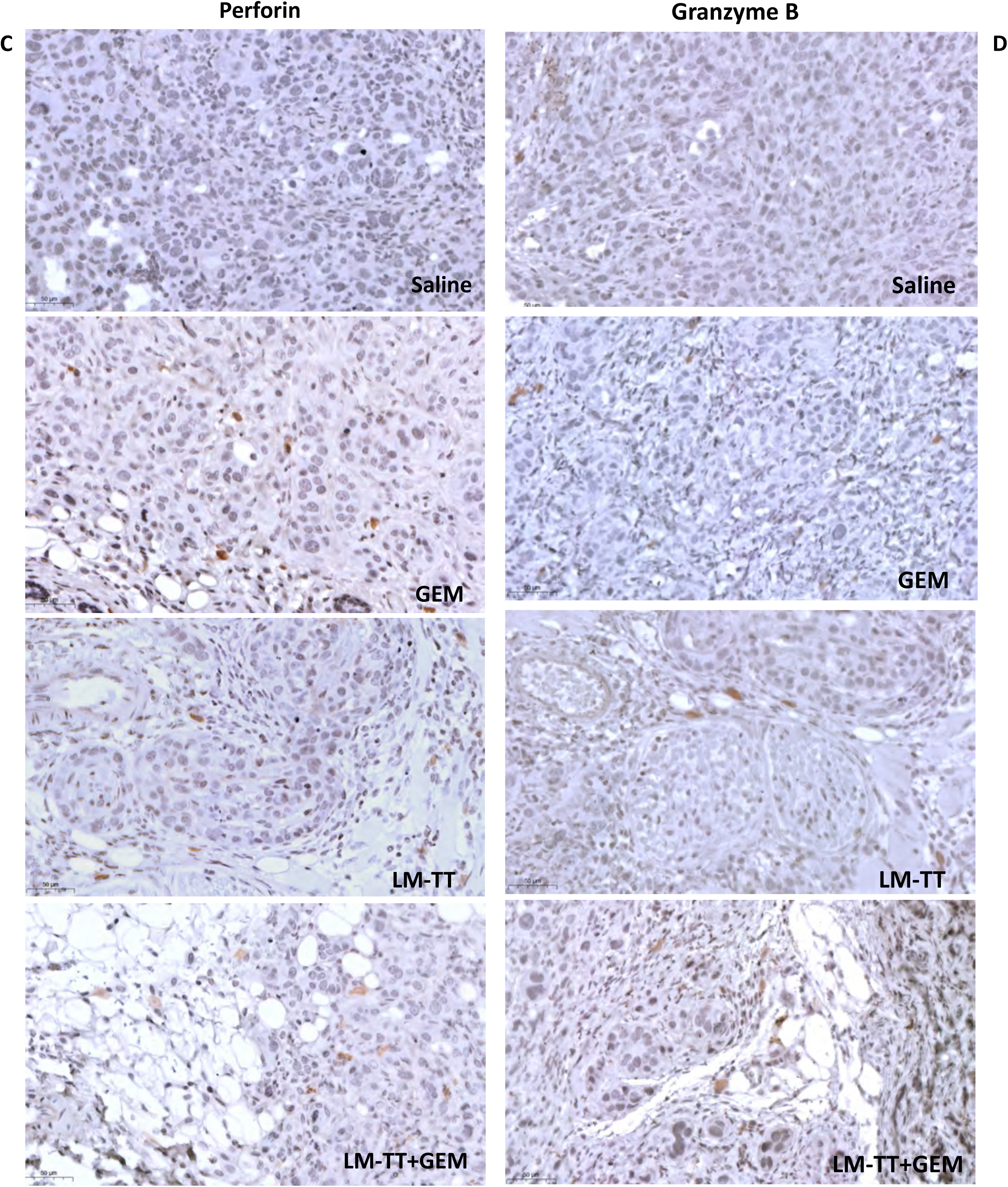
Perforin production and Granzyme B production in ovary tumors. Id8p53-/-Luc mice (ovary model) received treatments with Listeria-TT+GEM, or control treatments, as outlined in Figure S1A. **CD,** Two days after the last treatment, tumor tissues were analyzed by IHC for the production of perforin and granzyme B. LM-TT = Listeria-TT.

**Table S1:** Minimal effect(s) of high* or low** doses of anti-PD1 antibodies on spleen, lungs, kidneys, liver and small intestine of C57BL/6 mice***.

|  | Group Low Dose |  |  | Group High Dose |  |  |
| --- | --- | --- | --- | --- | --- | --- |
|  | Animal 1 | Animal 2 | Animal 3 | Animal 4 | Animal 5 | Animal 6 |
|  | Slides 1-2 | Slides 3-4 | Slides 5-6 | Slides 7-8 | Slides 9-10 | Slides 11-12 |
| **Spleen**<br>Increased extramedullary hematopoiesis | Minimal (1) | Minimal (1) | Minimal (1) | Slight (2) | Minimal (1) | Minimal (1) |
| **Lungs**<br>Hemorrhage, acute, focal<br>Inflammation, interstitial<br>Granuloma, focal | -<br>-<br>- | Minimal (1)<br>-<br>- | Minimal (1)<br>-<br>- | Minimal (1)<br>-<br>Minimal (1) | Slight (2)<br>Minimal (1)<br>- | Minimal (1)<br>-<br>- |
| **Kidneys (HE & PAS)**<br>Perirenal fat, infiltration lymphocytes | - | - | - | - | Minimal (1) | - |
| **Liver**<br>Extramedullary hematopoiesis<br>Necrosis, inflammation<br>Inflammation, focal | -<br>-<br>- | -<br>-<br>- | -<br>Minimal (1)<br>- | Minimal (1)<br>-<br>- | -<br>-<br>- | -<br>-<br>Minimal (1) |
| **Small Intestine**<br>Serosa, granuloma | - | - | Minimal (1) | - | - | - |

\*high dose of anti-PD1 antibodies (130 µg/dose)
\*\* low dose of anti-PD1 (50 µg/dose)
\*\*\* C57BL/6 mice of 6-8 weeks without cancer were treated with 4 high or low doses of anti-PD1 antibodies, each 4 days apart, as indicated in Supplemental Fig 3B. At the end of treatment, mice were euthanized and organs were subjected to pathological analysis to evaluate potential toxic effects of anti-PD1 antibodies.

## KEY RESOURCES TABLE

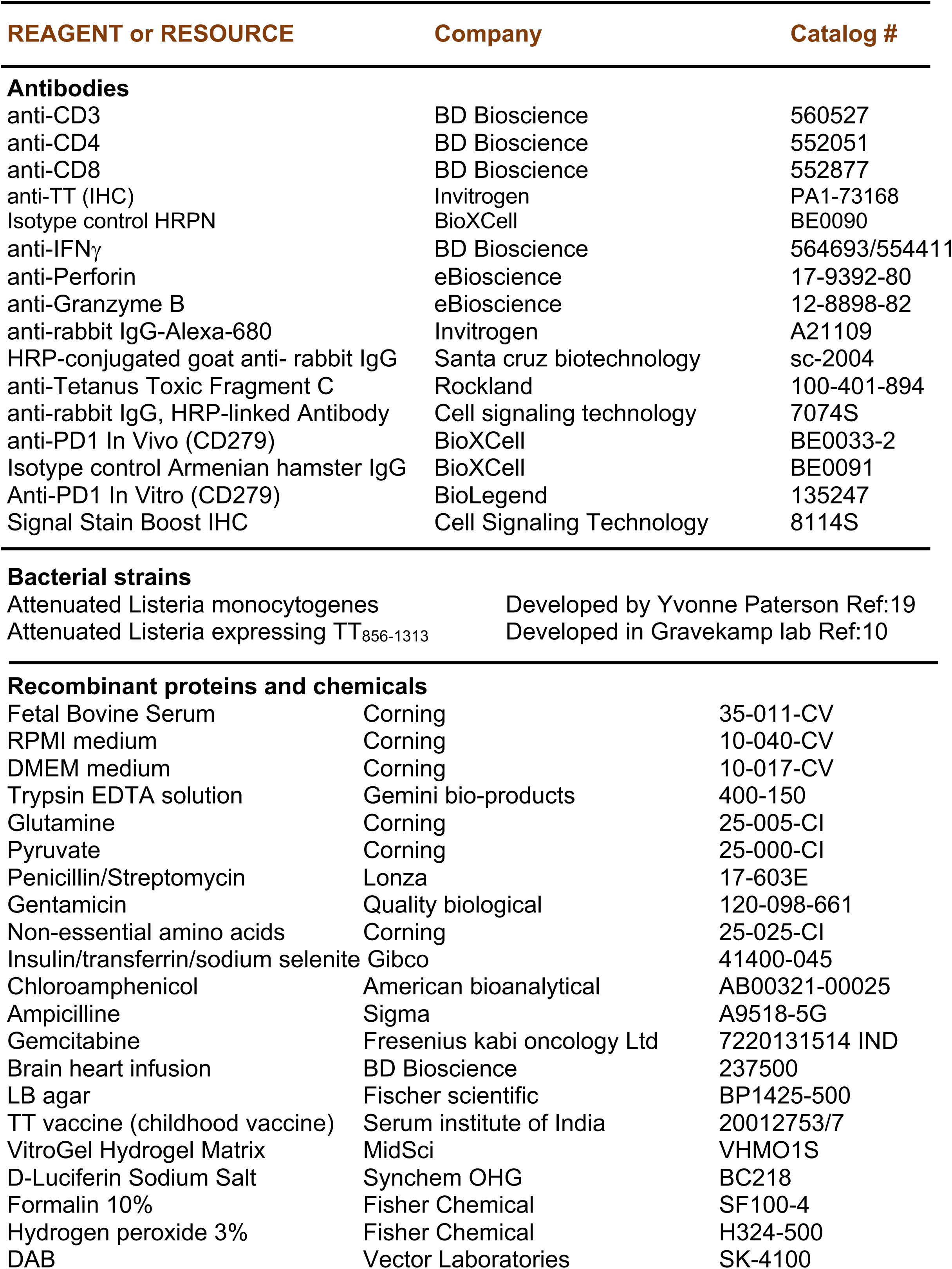
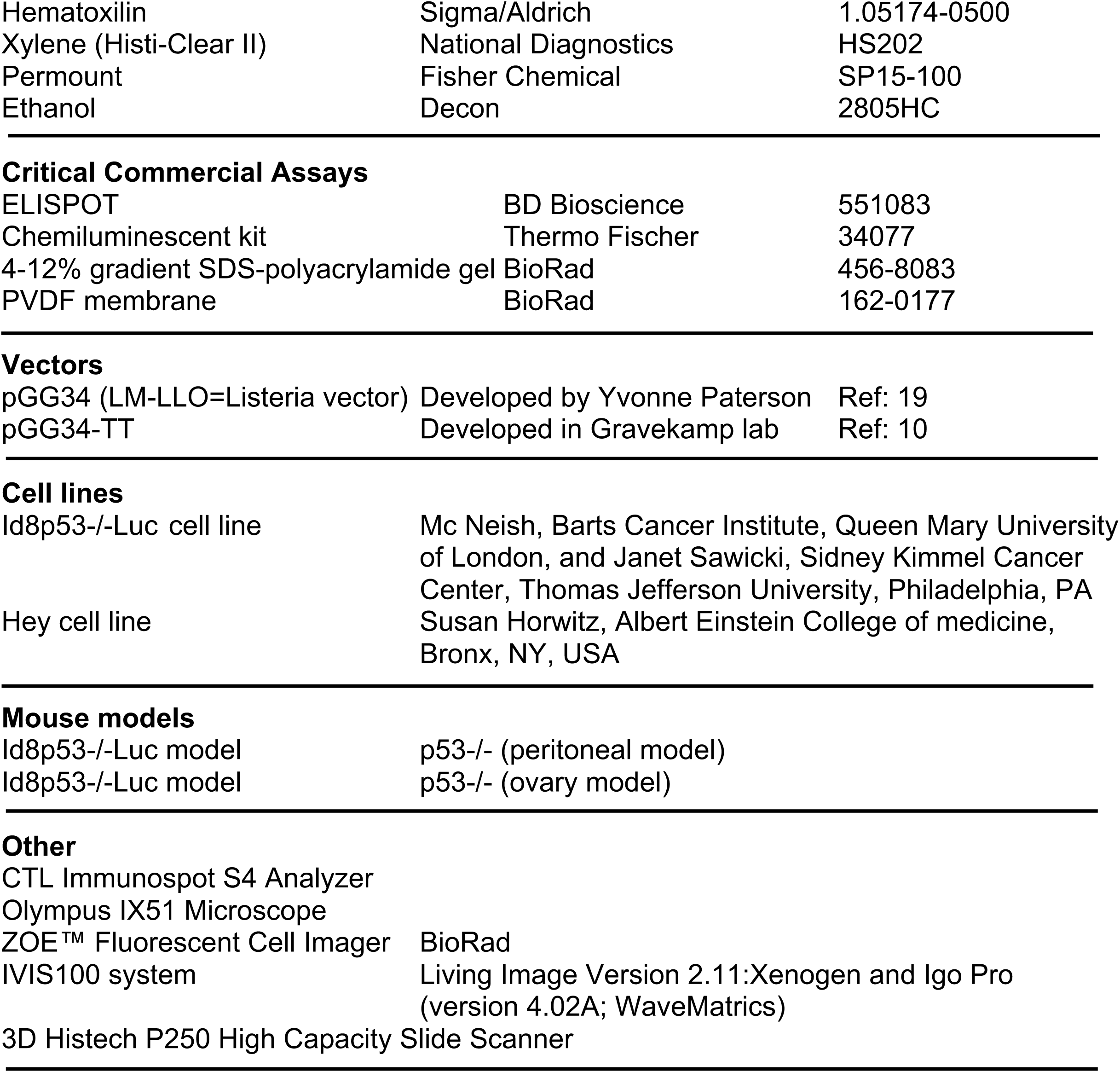

## Notes

### Competing Interest Statement

C.G. is the inventor of the Listeria-Recall Antigen Technology, which was developed in her laboratory, and is described in a patent application 11213577 (granted in the United States, China and Japan). The patent is licensed to Loki Therapeutics. C.G. is a stockholder of Loki Therapeutics and is an employee of Albert Einstein College of Medicine. All other authors declare that they have no competing interests.

### Summary of Updates

Expression profiles and biologic pathways have been analyzed in the ovarian tumors to study why the low dose of anti-PD1 was more effective than the high dose anti-PD1 antibodies, when combined with Listeria-TT and Gemcitabine in mice with ovarian cancer. This was added to the manuscript.

